# Targeting the FBXL12–FANCD2 Pathway Disrupts Replication Stress Tolerance in MYCN-Driven Neuroblastoma

**DOI:** 10.64898/2026.08.29.745966

**Authors:** Jinjiang Chou, Alena Malyukova, Alessandro Simone Bordonaro, Krzysztof Dygon, Lena Litzenburger, Ervelina Dalani, Jiayu Xiao, Conny Tümmler, Georgios Mermelekas, Janith Seniveratne, Magdalena Paolino, Juha Rantala, Lukas M Orre, Glenn Marshall, John Inge Johnsen, Malin Wickström, Andrä Brunner, Olle Sangfelt

## Abstract

MYCN amplification drives replication stress in high-risk neuroblastoma, yet how MYCN-amplified tumour cells tolerate this stress to sustain proliferation remains poorly understood. Here we show that FBXL12, an SCF ubiquitin ligase substrate receptor that targets the Fanconi anaemia protein FANCD2 for degradation at replication forks, as well as the broader Fanconi anaemia and replication stress transcriptional program are elevated in high-risk and MYCN-amplified neuroblastoma. High FBXL12 expression independently predicts poor survival across neuroblastoma patient cohorts. FBXL12 loss stabilizes FANCD2 on chromatin, elevates ATR-dependent replication stress signalling and DNA damage during S phase, and impairs proliferation of MYCN-amplified neuroblastoma cells in vitro and in vivo. Mechanistically, MYCN directly engages the FBXL12-FANCD2 complex and antagonises FBXL12-mediated degradation of FANCD2 at replication forks, revealing that the oncogenic driver of replication stress also actively preserves the chromatin-bound FANCD2 pool required to tolerate it. Beyond S phase, FBXL12 loss disrupts FANCD2-dependent mitotic DNA synthesis and transmits unresolved replication intermediates into daughter cells. FBXL12-deficient cells consequently show transcriptional activation of MYC target gene, ATR, and mTOR signalling programs, and this pathway-concordant state confers differential sensitivity to ATR, and mTOR-targeting compounds, nominating candidate therapeutic strategies for this disease subset. Together, these findings define a MYCN-FBXL12-FANCD2 axis as a clinically relevant vulnerability in high-risk neuroblastoma.

**HIGHLIGHTS:**

- FBXL12 is upregulated in high-risk neuroblastoma and predicts poor survival
- FBXL12 loss traps FANCD2 on chromatin and drives S-phase DNA damage
- MYCN antagonises FBXL12-mediated degradation of FANCD2 at replication forks
- FBXL12 loss impairs mitotic DNA synthesis and damages daughter cells
- FBXL12-deficient cells show ATR/mTOR activation and drug sensitivity

## INTRODUCTION

Neuroblastoma is the most common extracranial solid tumour of childhood, arising from undifferentiated sympathetic neural crest progenitors. Amplification of the MYCN oncogene occurs in approximately 20% of cases overall and in up to 40% of high-risk tumours, and remains the single genetic alteration most consistently linked to aggressive disease, treatment resistance and poor survival (Tonini et al., 1997, Campbell et al., 2017). Despite intensive multimodal therapy, long-term survival among high-risk patients remains below 50%, underscoring the need to define the molecular dependencies that allow MYCN-amplified tumour cells to sustain their proliferative programme and to identify how these dependencies might be therapeutically exploited.

Oncogenic MYCN activity drives replication stress. Aberrant origin firing, transcription-replication conflicts and nucleotide pool imbalance slow or stall replication forks and generate DNA damage during S phase (Halazonetis et al., 2008, Kotsantis et al., 2018). MYC family proteins contribute to this process directly. c-MYC binds the pre-replicative complex and increases the density of active origins independently of its transcriptional activity, and this non-transcriptional activity requires the CMG helicase component CDC45 (Dominguez-Sola et al., 2007, Srinivasan et al., 2013). In neuroblastoma specifically, MYCN expression reduces replication fork speed and increases fork stalling, sensitising cells to agents that further compromise fork stability, such as PARP inhibitors (King et al., 2020). Consistent with a broader dependency on the replication-stress response in MYC-driven disease, Myc-induced lymphomas require ATR and CHK1 for their development and survival, and MYC overexpression sensitises multiple myeloma cells to ATR inhibition (Murga et al., 2011, Cottini et al., 2014). Oncogene-induced replication stress is not, however, uniformly tumour-suppressive. A subset of stressed cells instead engages compensatory pathways that permit continued proliferation despite chronic fork instability, a phenomenon termed replication stress tolerance (Igarashi et al., 2024). This tolerance is frequently a specific, druggable property of the tumour cell rather than of normal tissue, making its molecular basis in MYCN-amplified neuroblastoma a rational route to new therapeutic targets.

The Fanconi anaemia (FA) pathway is a key mediator of replication stress tolerance. Upon fork stalling, the FANCD2-FANCI (ID2) heterodimer is recruited to and clamps around chromatin, where it protects nascent DNA, licenses fork remodelling and facilitates new origin firing (Schlacher et al., 2012, Lossaint et al., 2013, Chen et al., 2015, Alcon et al., 2020). This protective function depends not only on FANCD2’s recruitment to chromatin but also on the timely termination of FA pathway activity once replication stress is resolved. FANCD2 is deubiquitinated by USP1-UAF1, a step required to switch off FA pathway signalling (Nijman et al., 2005, Oestergaard et al., 2007). We recently identified FBXL12-mediated ubiquitination and proteasomal degradation as an additional route for clearing chromatin-associated FANCD2 during cyclin E-driven replication stress, and upon FBXL12 loss, FANCD2 itself becomes a barrier to fork progression (Brunner et al., 2023).

Whether an analogous turnover mechanism operates, and is regulated by the driving oncogene itself in other contexts of high replication stress has not been established. This question is particularly pertinent to MYCN-amplified neuroblastoma, where MYC-family proteins engage the replication-stress response directly. MYC can multimerise at stalled forks to physically shield them and stabilise fork-associated FANCD2 in a CHK1-and FANCI-dependent manner (Solvie et al., 2022), and MYCN additionally recruits AURKA-A to chromatin to support S-phase progression (Otto et al., 2009, Richards et al., 2016). Consistent with a broader, non-transcriptional role for MYC-family proteins in the genomic stress response, MYCN has been shown to adopt a distinct, non-transcriptional physical state under S-phase and transcriptional stress (Papadopoulos et al., 2022, Papadopoulos et al., 2024), Moreover, MYC has recently been shown to physically associate with FANCD2-marked chromatin in a manner that intensifies under replication stress (Cohn et al., 2026). These precedents suggest that MYCN does not simply generate replication stress as a by-product of proliferation but actively shapes how the FA pathway is engaged and resolved at stressed forks. This engagement may extend beyond the completion of S phase into the processing of under-replicated DNA in mitosis, via mitotic DNA synthesis (MiDAS) and the mitotic surveillance functions of the FA pathway (Naim and Rosselli, 2009, Minocherhomji et al., 2015, Bhowmick et al., 2016).

Here, we show that FBXL12 and the broader Fanconi anaemia pathway are elevated in high-risk and MYCN-amplified neuroblastoma, where high FBXL12 expression independently predicts poor survival. We define a MYCN-FBXL12-FANCD2 axis in which MYCN antagonises FBXL12-mediated FANCD2 degradation at replication forks and show that disrupting this axis impairs both S-phase fork progression and mitotic DNA synthesis, transmitting unresolved replication intermediates into daughter cells. Rather than simply halting proliferation, chronic disruption selects for an adaptive transcriptional and drug-sensitivity state that is pathway-concordant with the underlying mechanism. Together, these findings identify regulated FANCD2 turnover as a replication-stress tolerance mechanism co-opted by MYCN, and establish the FBXL12-FANCD2 axis as a clinically relevant vulnerability that confers differential sensitivity to ATR, and mTOR-targeting agents in high-risk neuroblastomas, warranting future exploration of combination strategies.

## RESULTS

### FBXL12 and the Fanconi anaemia pathway are upregulated in high-risk neuroblastoma and predict adverse clinical outcome (Figure 1, Figure S1)

To determine whether replication stress pathway activity distinguishes neuroblastoma risk groups, we scored a validated replication stress gene-expression signature (Takahashi et al., 2022) in an annotated multi-cohort neuroblastoma expression dataset assembled by Cangelosi et al. (Cangelosi et al., 2020). High-risk tumours had significantly higher replication stress signature scores than low-risk tumours (Figure 1A). Similarly, MYCN-amplified tumours had higher scores than non-amplified tumours (Figure 1B). Pathway enrichment analysis comparing high-risk and low-risk neuroblastoma transcriptomes identified significant enrichment of DNA replication, ribosome biogenesis, RNA transport, and the Fanconi anaemia pathway (Figure 1C). To assess whether expression of Fanconi anaemia pathway components and associated genes was linked to clinical outcome, we performed 23 separate multivariable Cox regression models, one per gene, each adjusted for age, MYCN amplification status, and stage 4 disease in the Kocak neuroblastoma cohort (Kocak et al., 2013). Expression of 11 of 23 analysed genes, was significantly associated with increased hazard of death (nominal p < 0.05), with FBXL12 showing the highest hazard ratio, identifying it as an independent adverse prognostic marker after adjustment for age, MYCN status, and stage (Figure 1D). For event-free survival, 16 of 23 genes were significantly associated with increased hazard of an event (Figure 1E). Repeating the analysis in the independent SEQC cohort (Zhang et al., 2015) supported a broader association between Fanconi anaemia pathway gene expression and adverse outcome in neuroblastoma (Figure S1A and S1B). Consistent with its association with adverse outcome, FBXL12 expression was also significantly elevated in high-risk disease across the larger multi-cohort dataset from (Cangelosi et al., 2020) (Figure 1F).

**Figure 1.**
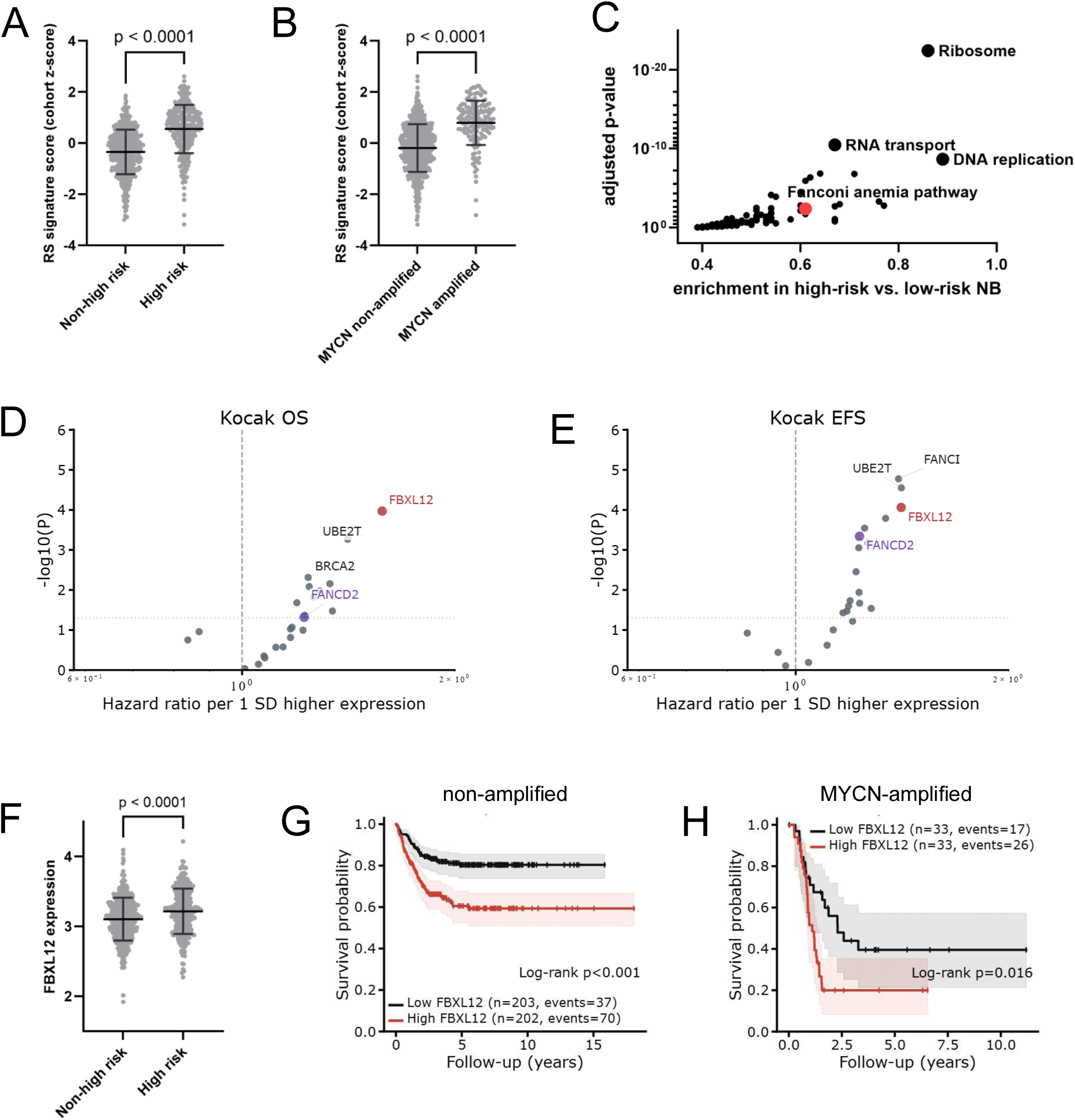
FBXL12 and the Fanconi anaemia pathway are upregulated in high-risk neuroblastoma and predict adverse outcome. (A) Replication-stress signature score in non-high-risk and high-risk neuroblastoma tumours. (B) Replication-stress signature score in MYCN non-amplified and MYCN-amplified neuroblastoma tumours. (C) Gene ontology and pathway enrichment analysis of transcripts enriched in high-risk compared with low-risk neuroblastoma. Selected pathways, including ribosome, RNA transport, DNA replication and Fanconi anaemia pathway, are highlighted. (D, E) Univariate Cox proportional-hazards analysis of Fanconi anaemia pathway genes in the Kocak cohort for overall survival (D) and event-free survival (E) Hazard ratios are shown per 1 SD higher gene expression. FBXL12 and FANCD2 are highlighted. (F) FBXL12 expression in non-high-risk and high-risk neuroblastoma tumours. (G, H) Kaplan-Meier survival analysis stratified by median FBXL12 expression in MYCN non-amplified (G) and MYCN-amplified (H) neuroblastoma subgroups. Differences between survival curves were evaluated using the log-rank test. Panels A–C and F use the Cangelosi cohort; panels D, E, G and H use the Kocak cohort.

Kaplan-Meier analysis revealed that high FBXL12 expression was associated with significantly reduced survival probability, in both MYCN-amplified and non-amplified subgroups (Figure 1G and 1H). Extended survival analyses confirmed poor event-free survival in FBXL12-high patients across the full Kocak (Figure S1C) and SEQC (Figure S1D) cohorts. Within the MYCN-amplified SEQC subgroup, FBXL12-high patients showed a trend toward inferior event-free survival that did not reach statistical significance (p = 0.186, Figure S1E), likely reflecting the limited size of this subgroup. The association was significant in the non-amplified subgroup (Figure S1F). High FANCD2 mRNA expression was similarly associated with poor event-free survival in MYCN non-amplified tumours (Figure S1G) but not in MYCN-amplified tumours (Figure S1H). MYCN mRNA levels correlated positively with FANCD2 mRNA levels across Kocak and SEQC patient cohorts (Figure S1I and S1J). This was further corroborated experimentally, as dox-induced MYCN depletion in the inducible SHEP model reduced FANCD2 mRNA levels (Figure S1K).

Together, these data show that a Fanconi anaemia and replication stress transcriptional programme, and FBXL12 specifically, are elevated in high-risk and MYCN-amplified neuroblastoma. High FBXL12 expression independently predicts poor clinical outcome, identifying the FBXL12-FANCD2 axis as a clinically relevant feature of aggressive disease.

### FBXL12 loss stabilises FANCD2, elevates S-phase damage, and impairs proliferation of MYCN-amplified neuroblastoma cells in vitro and in vivo (Figure 2, Figure S2)

To directly interrogate FBXL12 function in MYCN-amplified neuroblastoma cells, we generated two independent FBXL12 knockout clones in SK-N-BE(2)C cells by CRISPR-Cas9 (Figure S2A, B). qPCR confirmed loss of FBXL12 transcript in both knockout clones, validating the knockouts at the mRNA level. FANCD2 and MYCN mRNA levels were largely unchanged in the FBXL12-KO cells (Figure S2C). Consistent with our previous findings in cyclin E-driven replication stress models (Brunner et al., 2023), FBXL12 depletion caused marked accumulation of FANCD2 protein in both knockout clones, without a corresponding increase in FANCD2 mRNA following either knockout or siRNA-mediated depletion of FBXL12 (Figure 2A, S2C), indicating that this stabilisation is post-transcriptional. FANCD2 accumulation was accompanied by elevated phospho-RPA32 Ser33, an ATR-dependent phosphorylation mark, and increased γH2AX (Figure 2A). This is consistent with accumulation of replication-associated DNA damage. FANCD2 stabilisation was similarly observed in Kelly FBXL12-knockout cells and following acute siRNA-mediated FBXL12 depletion in Kelly and IMR-32 cells, demonstrating that FBXL12-dependent control of FANCD2 is conserved across MYCN-amplified neuroblastoma cell lines (Figure S2D).

**Figure 2.**
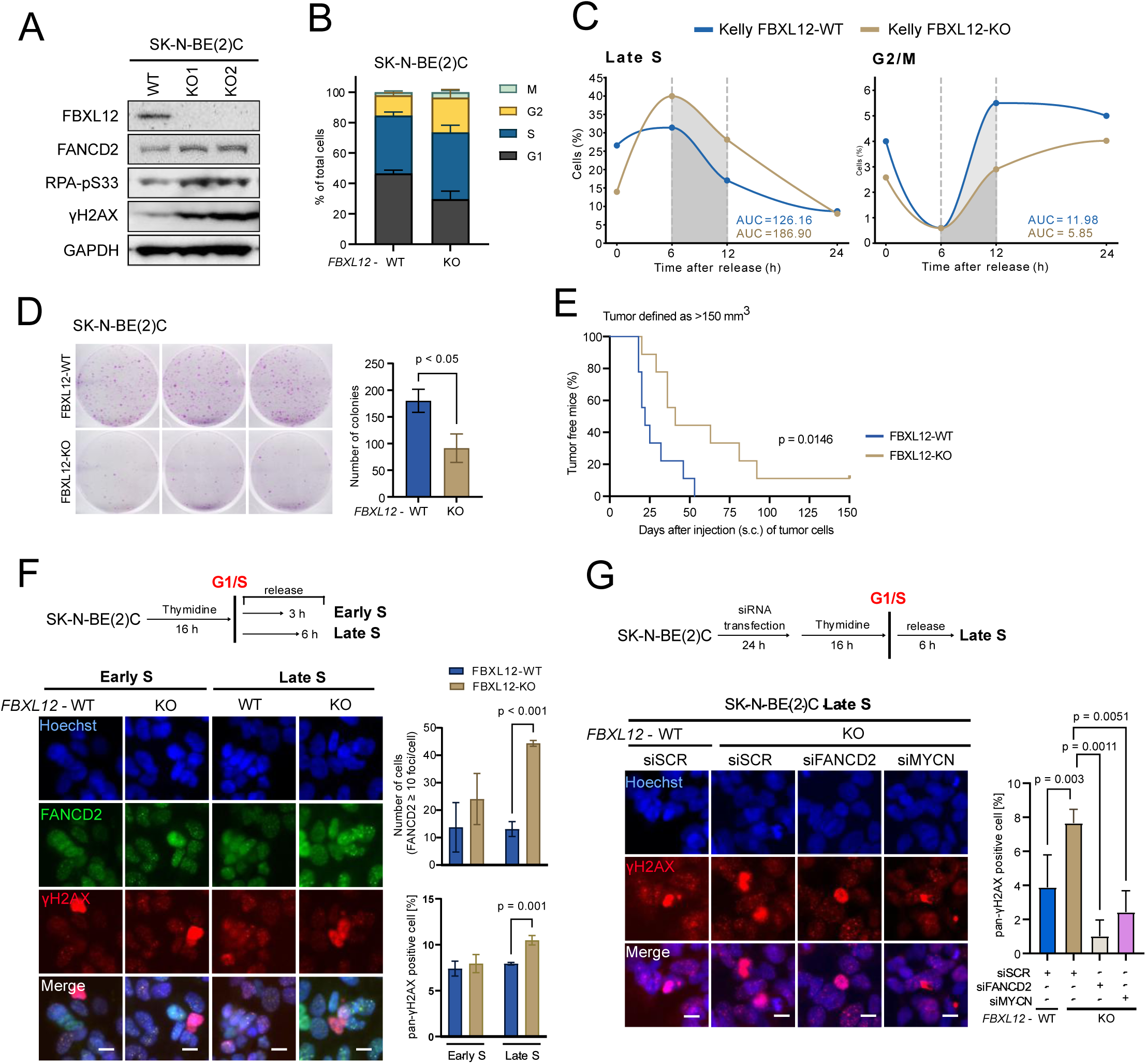
FBXL12 loss stabilises FANCD2, elevates S-phase damage, and impairs proliferation. (A) Immunoblots of FANCD2, pS33-RPA32, and γH2AX in SK-N-BE(2)C wild-type and two FBXL12-knockout clones. (B) Cell-cycle distribution evaluated by high content imaging staining of EdU in SK-N-BE(2)C wild type (WT) and FBXL12-knockout (KO) clone. (C) Kelly WT and FBXL12-KO cells were pulse-labelled with EdU for 30 min and released for the indicated times. EdU incorporation and DNA content (Hoechst, NN) were analysed by flow cytometry. The percentages of EdU-positive cells in late S phase and 4N cells in G2/M are shown over time. AUC was calculated over the 6 −12 h interval (shown in grey). (D) Colony formation of SK-N-BE(2)C WT and FBXL12-KO clones. (E) Kaplan-Meier curves of the percentage of tumour-free mice following subcutaneous (s.c) injection of SK-N-BE(2)C WT (blue, n = 9) and FBXL12-KO (light brown, n = 9) clones. Tumour formation was defined as a tumour volume greater than 150 mm^3^. P-value of Logrank (Mantel-Cox) test is presented. (F) FANCD2 foci and pan-γH2AX quantification in SK-N-BE(2)C WT and FBXL12-KO clones at 3 (early S phase) and 6 h (late S phase) post-release from 16 h thymidine (2 mM) block, scale bar: 10 µm. (G) Pan-γH2AX quantification at 6 h post-release (late S phase) from thymidine in SK-N-BE(2)C WT and FBXL12-KO clones with/without FANCD2 or MYCN co-depletion, scale bar: 10 µm. (B, D, F and G) Data shown are means with error bars indicating SD.

High-content imaging of EdU-pulsed SK-N-BE(2)C cells showed an altered cell-cycle distribution in FBXL12-knockout cells, with a relative decrease in G1 and accumulation in S phase (Figure 2B). To directly test this, we pulse-labelled Kelly and SK-N-BE(2)C wild-type and FBXL12-knockout cells with EdU and tracked the labelled cohort by flow cytometry over 0–24 h following release. By 10–12 h, wild-type cells had largely cleared late S phase and entered G2/M, whereas FBXL12-knockout cells remained enriched in late S with delayed G2/M entry (Figure 2C and Figure S2E), confirming a significant delay in S-phase progression consistent with elevated replication stress and slowed fork progression. Colony formation assays with early passage showed that SK-N-BE(2)C FBXL12-knockout cells formed approximately 50% fewer colonies than wild-type cells when seeded at equal density, indicating that FBXL12 loss impairs proliferative capacity without causing complete growth arrest (Figure 2D).

To determine whether FBXL12 loss affects tumour growth in vivo, wild-type and FBXL12-knockout SK-N-BE(2)C cells were injected subcutaneously into NMRI nu/nu mice across two independent experiments (pilot, n=3/group; follow-up, n=6/group; combined n = 9 /group). Tumours in the FBXL12-knockout group took significantly longer to reach 150 mm^3^ than wild-type tumours in the pooled cohort (log-rank p = 0.0146, Figure 2E). Median time to tumour establishment was 25 days in wild-type tumours, with delayed onset in knockout tumours. This difference was robust across multiple size thresholds (Figure S2F) and remained significant when accounting for between-experiment variation (p = 0.042, see Figure legend and methods). Western blot analysis of excised tumours confirmed loss of FBXL12 and elevated γH2AX in knockout tumours relative to wild-type controls (Figure S5B).

To characterise FANCD2 chromatin dynamics across S phase in the absence of FBXL12, SK-N-BE(2)C wild-type and knockout cells were synchronised by thymidine block and released into S phase. FANCD2 nuclear foci progressively accumulated in knockout cells relative to wild-type at 3 and 6 hours post-release, reaching statistical significance at 6 hours (Figure 2F), consistent with aberrant FANCD2 trapping on chromatin during S-phase progression. At 6 hours, FBXL12-knockout cells showed significantly elevated γH2AX foci relative to wild-type (Figure 2G). Co-depletion of FANCD2 by siRNA rescued the γH2AX increase in knockout cells, indicating that S-phase DNA damage upon FBXL12 loss is FANCD2-dependent (Figure 2G). The FANCD2-dependence of this phenotype was confirmed using acute siRNA-mediated FBXL12 depletion with FANCD2 co-depletion rescue in IMR-32 and Kelly cells (Figure S2G), supporting conservation across MYCN-amplified lines. Depletion of MYCN using siRNA also rescued the elevated γH2AX in FBXL12-knockout cells (Figure 2G), showing that MYCN itself contributes to the S-phase damage phenotype rather than being a passive bystander of FBXL12 loss. Since FBXL12 is already absent in these cells, this rescue cannot reflect restored FBXL12-mediated FANCD2 turnover. It instead points to a separate, FBXL12-independent role for MYCN, addressed below (Figure 3).

**Figure 3.**
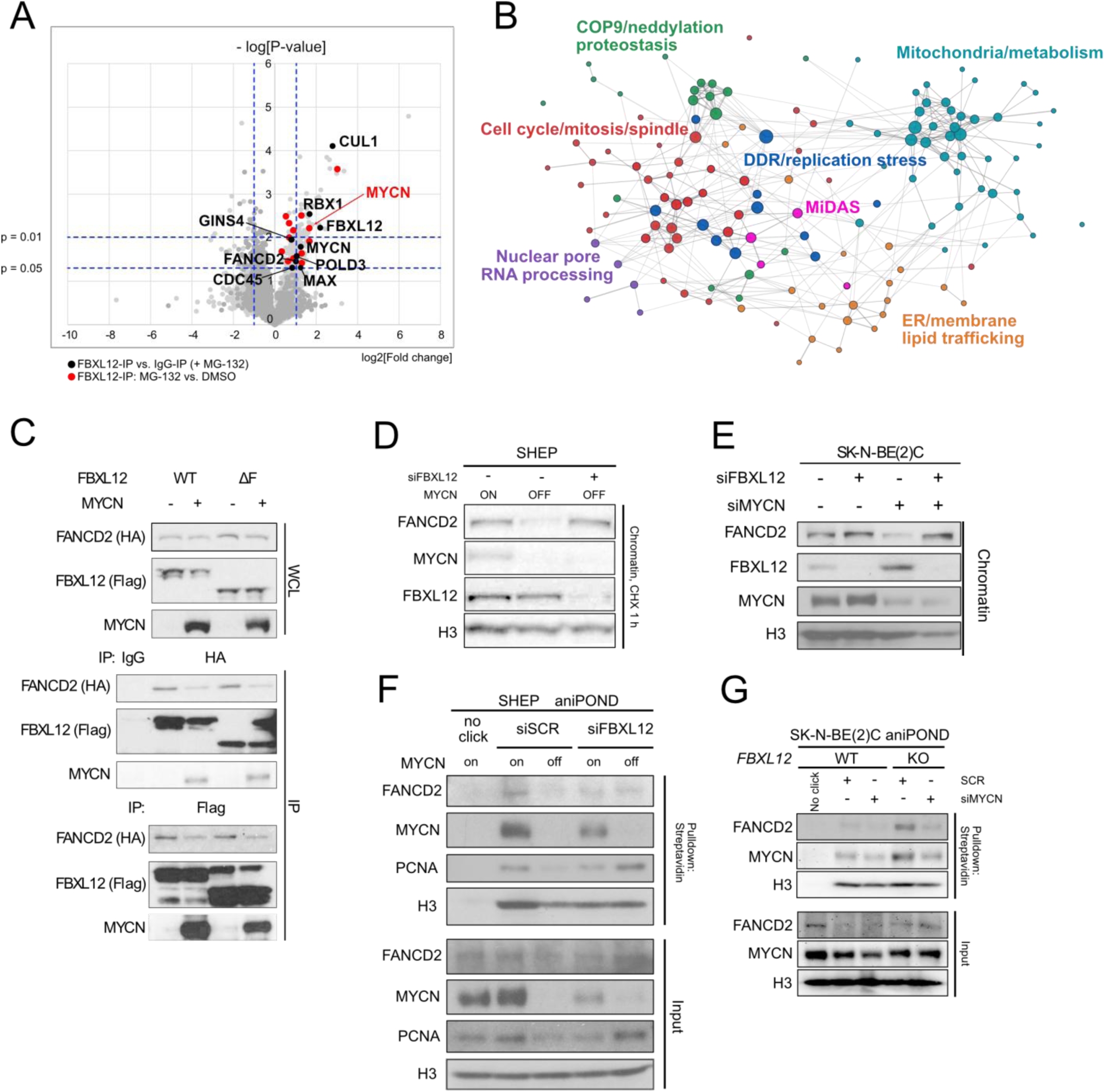
MYCN antagonises FBXL12-mediated degradation of FANCD2 at replication forks. (A) Volcano plot of FBXL12-interacting proteins identified by FBXL12 IP-mass spectrometry in SK-N-BE(2)C cells treated with vehicle or MG132. Dark gray points indicate proteins enriched in FBXL12-IP compared with IgG-IP under MG132 treatment. Light gray points indicate proteins enriched in the MG132-treated group compared with the DMSO-treated group in FBXL12-IP samples. Black and red points highlight SCF complex components, MYCN/MAX and FBXL12/FANCD2. Horizontal dashed lines indicate significance thresholds of p = 0.05 and p = 0.01, and vertical dashed lines indicate a log₂ fold-change cutoff of ±1. (B) STRING network of enriched proteins grouped by unbiased functional modules. Node size indicates STRING degree; colours mark functional clusters, with DDR, replication stress/MiDAS, and mitotic proteins highlighted. (C) Immunoprecipitation of FANCD2-HA from HEK293T FBXL12-KO cell chromatin lysates transfected with the indicated constructs. (D) Immunoblot of FANCD2 in chromatin fractions isolated from SHEP cells with different MYCN expression status and transfected with control or FBXL12 siRNA, following a 1-h CHX chase. (E) Immunoblot of FANCD2 in chromatin fractions isolated from SK-N-BE(2)C cells following MYCN and/or FBXL12 siRNA transfection. (F) aniPOND analysis of MYCN, FANCD2 and PCNA at newly replciated DNA in SHEP cells with different MYCN expression status and siFBXL12 transfection. (G) aniPOND analysis of MYCN and FANCD2 in SK-N-BE(2)C WT and FBXL12-KO cells with/without MYCN siRNA transfection.

Together, these data show that FBXL12 restrains FANCD2 chromatin accumulation during S phase. Its loss drives FANCD2-dependent replication stress and DNA damage that impairs the proliferation of MYCN-amplified neuroblastoma cells both in vitro and in vivo.

### MYCN antagonises FBXL12-mediated degradation of FANCD2 at replication forks (Figure 3, Figure S3)

To identify proteins associated with FBXL12 in MYCN-amplified neuroblastoma cells, we immunopurified endogenous FBXL12 from Kelly cell lysates under basal conditions and following proteasome inhibition with MG132, and analysed co-purifying proteins by mass spectrometry. FBXL12 immunoprecipitates were significantly enriched, relative to IgG controls, for proteins spanning several functional categories, including the SCF complex components CUL1 and RBX1, the COP9 signalosome, the replication fork factors CDC45 and GINS1/4, the DNA repair factors FANCD2 and POLD3, and both MYCN and its obligate dimerisation partner MAX (Figure 3A and table S1). Gene ontology enrichment analysis of the FBXL12 interactome confirmed significant enrichment of cell cycle, mitosis, mitosis DNA synthesis (MiDAS), DNA damage repair (DDR), nuclear pore RNA processing, mitochondria/metabolism and neddylation proteostasis terms (Figure 3B, Figure S3A).

MG132 treatment substantially increased MYCN enrichment in FBXL12 immunoprecipitates, with MYCN ranking among the top three most enriched proteins under proteasome-inhibited conditions (Figure 3A, table S1), indicating that the FBXL12-MYCN interaction increases as MYCN accumulates. The FBXL12-MYCN interaction was validated by co-immunoprecipitation in Kelly, where MYCN co-purified weakly with FBXL12 under basal conditions but was markedly enriched after MG132 treatment, accompanied by co-purification of FANCD2 (Figure S3B). To test whether MYCN modulates the FBXL12–FANCD2 interaction, HEK293 cells were co-transfected with FBXL12-WT or ΔF, a construct lacking the F-box domain required for SCF assembly and thus degradation activity, FANCD2-HA, and MYCN, in the presence of MG132 (Figure 3C). Co-IP in both directions showed reduced FANCD2–FBXL12 association upon MYCN overexpression, most clearly in the Flag-IP (FBXL12 pulldown), where reduced FANCD2 co-purified despite comparable FBXL12 recovery, in both the WT and ΔF backgrounds, indicating this reduction is independent of the FBXL12 F-box domain. MYCN itself co-precipitated with FBXL12 only upon MYCN transfection, confirming the interaction. Further supporting an interplay among FANCD2, FBXL12 and MYCN in neuroblastoma cells, immunoprecipitation of endogenous FANCD2 from SHEP cells also recovered both FBXL12 and MYCN (Figure S3C). Taken together these data indicate that MYCN antagonises the FBXL12-FANCD2 interaction and reduces FANCD2 binding to the SCF-FBXL12 complex.This aligns with emerging noncanonical roles for MYC family proteins, including a compensatory fork-protection response mediated by MYC/MYCN multimerization at stalled forks under replication-stress (Solvie et al., 2022).

Given that FBXL12-dependent turnover of chromatin-bound FANCD2 appeared central to limiting replication stress (Figure 2), we asked whether MYCN itself regulates this turnover. In SHEP neuroblastoma cells with doxycycline-regulated MYCN expression, chromatin fractionation after cycloheximide treatment showed that FANCD2 was markedly reduced on chromatin upon MYCN depletion, despite only a modest reduction in FBXL12 protein levels (Figure 3D). siRNA-mediated FBXL12 co-depletion fully rescued FANCD2 chromatin association in MYCN-depleted cells (Figure 3D), indicating that FANCD2 instability upon MYCN loss is mediated by FBXL12-dependent proteasomal degradation. Concordant results were obtained in SK-N-BE(2)C cells. siRNA-mediated MYCN knockdown reduced chromatin-bound FANCD2, and co-depletion of FBXL12 fully rescued FANCD2 chromatin association (Figure 3E).

To test whether MYCN and FANCD2 co-occupy replication forks, and whether FBXL12 controls their fork association, we used isolation of proteins on nascent DNA in SHEP cells under MYCN-on and MYCN-off conditions, with or without FBXL12 knockdown. MYCN and FANCD2 were both present at forks in MYCN-on cells, together with the fork marker PCNA (Figure 3F). MYCN depletion reduced FANCD2 and PCNA at forks, consistent with impaired fork progression, and FBXL12 co-depletion fully restored FANCD2 fork association and PCNA levels in MYCN-off cells (Figure 3F). The same relationship held in SK-N-BE(2)C FBXL12-knockout cells following siMYCN treatment (Figure 3G), loss of FANCD2 fork association upon MYCN depletion was largely dependent on FBXL12, with only a modest residual reduction in FBXL12-KO cells. Together, these data demonstrates that FBXL12 is epistatic to MYCN in controlling FANCD2 fork occupancy, placing MYCN upstream of FBXL12-mediated FANCD2 turnover at replication forks. However, this fork-occupancy epistasis cannot fully account for the MYCN-dependent rescue of S-phase damage in FBXL12-KO cells (Figure 2G). Since FBXL12-mediated FANCD2 degradation is already absent in these cells, MYCN depletion cannot act through this route, and the residual, FBXL12-independent reduction in fork-associated FANCD2 may instead contribute to the S-phase damage rescue.

Together, these data suggest that MYCN antagonises FBXL12-mediated FANCD2 degradation at replication forks, maintaining a balanced level of chromatin-associated FANCD2 that supports replication stress tolerance under MYCN-driven proliferative stress. This balance is disrupted by loss of FBXL12-mediated turnover, which instead traps FANCD2 on chromatin and triggers excessive replication stress.

### FBXL12 loss impairs mitotic DNA synthesis and drives FANCD2-dependent DNA damage transmission to daughter cells (Figure 4, Figure S4)

Mass spectrometry analysis of the FBXL12 interactome identified several proteins exclusively in FBXL12 immunoprecipitates and absent from IgG controls (Figure S4A). These included ATR and GINS1, placing FBXL12 at active replication forks in proximity to the ATR signalling machinery, as well as multiple mitotic regulators, including the centrosome components CEP192 and CEP170, the kinetochore factor KNL1, and NSMCE4A, a subunit of the SMC5/6 complex required for resolving replication intermediates that persist into mitosis (Menolfi et al., 2015). This suggested that FBXL12 function extends beyond S phase into mitotic genome maintenance. To test this directly, we examined mitotic DNA synthesis, mitotic DNA damage, and the transmission of unresolved replication intermediates into the subsequent G1 phase in FBXL12-deficient cells.

**Figure 4.**
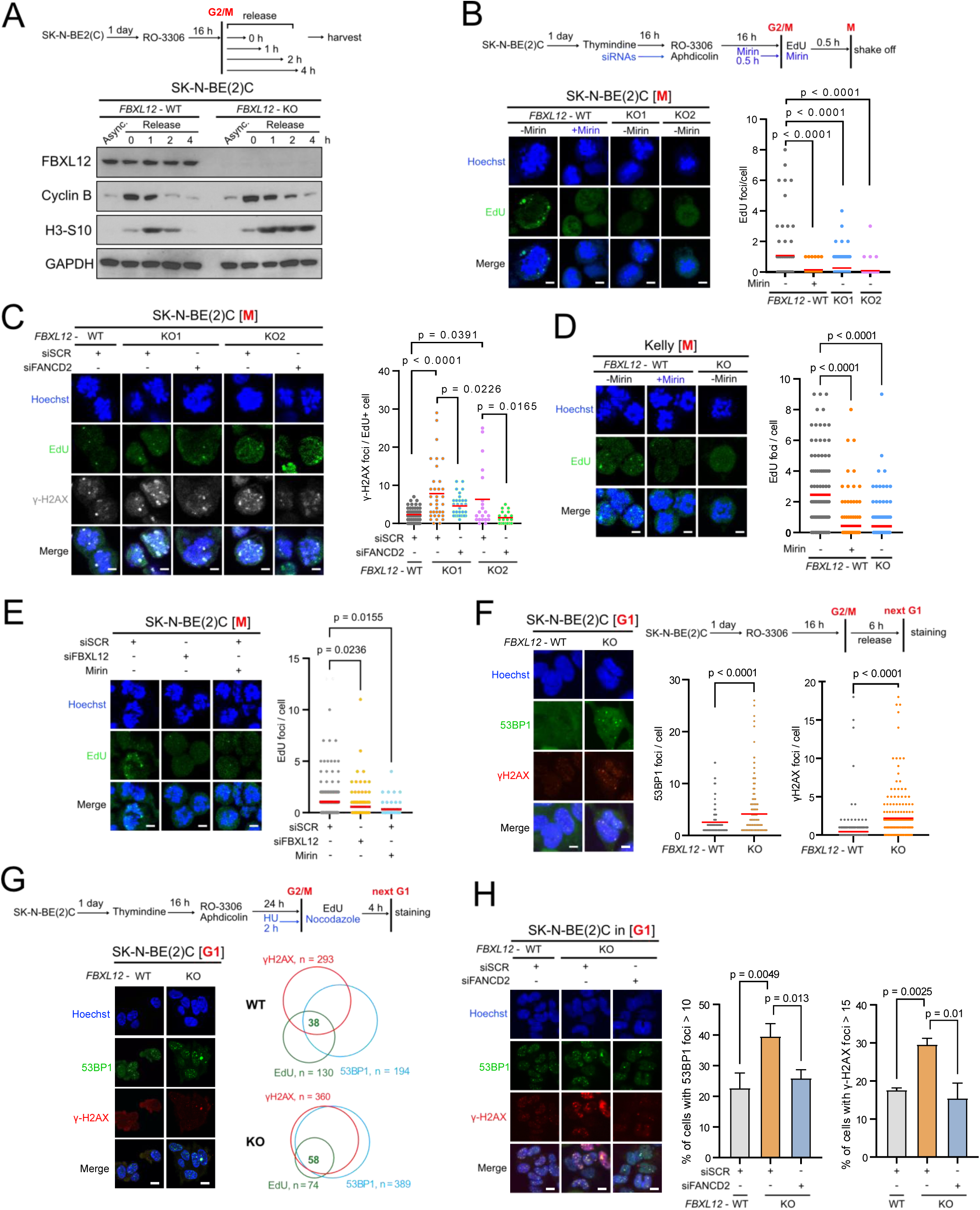
FBXL12 loss impairs mitotic DNA synthesis and drives FANCD2-dependent damage transmission. (A) Immunoblots of Cyclin B and pS10-H3 in SK-N-BE(2)C WT and FBXL12-KO clones released from RO-3306 (9 µM) synchronization. (B) SK-N-BE(2)C WT and FBXL12-KO cells were synchronized as indicated in the experimental scheme, pulse-labelled with EdU and released into the prometaphase. MiDAS was evaluated by counting EdU foci in SK-N-BE(2)C WT and FBXL12-KO prometaphase cells. Mirin (10 μM) treatment served as a positive control. Scale bar: 5 μm. (C) Quantification of prometaphase γH2AX foci in SK-N-BE(2)C WT and FBXL12-KO clones with/without FANCD2 co-depletion. Scale bar: 5 μm. (D) Quantification of MiDAS in Kelly WT and FBXL12-KO cells in prometaphase. Mirin (10 μM) treatment served as a positive control. (E) Quantification of MiDAS in SK-N-BE(2)C cells upon short-term FBXL12 loss by FBXL12 siRNA transfection for 48 h. Scale bar: 5 μm. (F) SK-N-BE(2)C WT and FBXL12-KO cells were synchronized as indicated in the experimental scheme, pulse-labelled with EdU and released into the next G1 phase. Representative images and quantification of 53BP1 and γH2AX in SK-N-BE(2)C WT and FBXL12-KO cells in next-G1 phase, scale bar: 5 μm. (G) SK-N-BE(2)C WT and FBXL12-KO cells were synchronized as indicated in the experimental scheme, pulse-labelled with EdU and released into the next G1 phase. Thirty G1-phase cells retaining EdU-positive signal from the initial labelling were analysed per condition. Venn diagrams show the overlap of residual EdU foci with 53BP1 and γH2AX foci. Numbers indicate the total number of foci analysed for each marker, with the number of EdU/53BP1/γH2AX triple-positive foci shown in the centre. (H) Quantification of 53BP1 and γH2AX foci in SK-N-BE(2)C WT and FBXL12-KO G1 daughter cells with/without FANCD2 co-depletion, scale bar: 10 μm. (B, C, D and E) Individual and mean values are plotted. (H) panel data shown are means with error bars indicating SD.

We first characterised mitotic progression in synchronised SK-N-BE(2)C wild-type and FBXL12-knockout cells released from CDK1 inhibition with RO-3306. In wild-type cells, cyclin B levels peaked at 0 to 1 hours post-release and declined progressively, and pS10-H3 peaked at 1 hour before rapidly diminishing by 2-to 4-hours, consistent with normal mitotic entry and exit (Figure 4A). FBXL12-knockout cells showed sustained cyclin B and pS10-H3 at 2 to 4 hours post-release, indicating delayed mitotic exit (Figure 4A), consistent with unresolved replication intermediates engaging a mitotic checkpoint response.

To directly assess mitotic DNA synthesis, SK-N-BE(2)C wild-type and FBXL12-knockout cells were synchronised with RO-3306 and aphidicolin, released into mitosis in the presence of EdU, and prometaphase cells were isolated by mitotic shake-off (Xu et al., 2021, Petropoulos et al., 2024, Barwacz et al., 2025). FBXL12-knockout cells showed a significant reduction of EdU-foci in prometaphase cells relative to wild-type (Figure 4B), demonstrating impaired mitotic DNA synthesis. The magnitude of this reduction was comparable to that achieved by MRE11 inhibition with Mirin in wild-type cells (Figure 4B). FANCD2 foci remained detectable in prometaphase cells of both genotypes (Figure S4B), indicating that the reduced mitotic DNA synthesis in FBXL12-knockout cells does not result from loss of chromatin-associated FANCD2, a condition previously reported to abolish mitotic DNA synthesis (Xu et al., 2021). Prometaphase γH2AX foci were significantly elevated in FBXL12-knockout cells (Figure 4C), and this increase was rescued by FANCD2 co-depletion (Figure 4C), demonstrating that mitotic DNA damage upon FBXL12 loss is FANCD2-dependent.

To determine whether impaired mitotic DNA synthesis is also observed beyond the SK-N-BE(2)C FBXL12-knockout clones, we extended the assay to Kelly cells and to acute siRNA-mediated FBXL12 knockdown in SK-N-BE(2)C cells. Both showed a significant reduction of EdU-foci in prometaphase cells relative to controls (Figure 4D, E), confirming that impaired mitotic DNA synthesis is a conserved feature of FBXL12 deficiency across MYCN-amplified neuroblastoma lines. To assess whether impaired mitotic DNA synthesis in FBXL12-knockout cells leads to transmission of unresolved damage into the next cell cycle, we used a previously described protocol (Lukas et al., 2011) in which synchronised cells were released into mitosis with EdU, collected by shake-off, and replated for analysis of the resulting G1 daughter cells. FBXL12-knockout daughter cells showed a significant increase in 53BP1 nuclear bodies relative to wild-type (Figure 4F), consistent with inheritance of under-replicated loci from the preceding mitosis. G1 γH2AX foci were similarly elevated in FBXL12-knockout cells (Figure 4F), indicating that unresolved replication intermediates persist as DNA damage beyond mitotic exit rather than being resolved prior to cell division. Venn diagram analysis of marker co-localisation showed that DNA lesions in knockout daughter cells arise predominantly at loci that failed to complete mitotic DNA synthesis, evidenced by disproportionate overlap between damage markers and residual EdU foci in knockout compared with wild-type cells (Figure 4G). Co-depletion of FANCD2 rescued the elevated 53BP1 and γH2AX signal in G1 daughter cells of FBXL12-knockout cells (Figure 4H), confirming that transmission of mitotic DNA damage is driven by aberrant FANCD2 stabilisation downstream of FBXL12 loss. siMYCN did not significantly rescue elevated G1 damage in FBXL12-knockout cells (Figure S4C), in contrast to its rescue of S-phase DNA damage (Figure 2G). This indicates that mitotic damage transmission is independent of MYCN activity, distinguishing the S-phase and mitotic arms of the FBXL12-dependent phenotype and pointing to FANCD2 dysregulation itself, rather than ongoing MYCN activity, as the driver of damage transmitted into G1.

Together, these data show that FBXL12 loss impairs mitotic DNA synthesis, elevates FANCD2-dependent mitotic DNA damage, and drives transmission of unresolved replication intermediates to daughter cells. The FBXL12-FANCD2 axis is required for faithful completion of DNA replication across both S phase and mitosis in MYCN-amplified neuroblastoma cells.

### FBXL12 loss and FANCD2 insufficiency converge on a common MYCN-linked stress-adaptation programme (Figure 5, Figure S5)

In parallel to the FBXL12-knockout model, we were unable to recover SK-N-BE(2)C clones with biallelic FANCD2 deletion, consistent with FANCD2 loss being poorly tolerated in this cell line. We instead obtained a monoallelic FANCD2-low clone (Figure 5A, S5A), to model a state of FANCD2 insufficiency and related phenotypes. In a separate, small xenograft cohort (n = 3), SK-N-BE(2)C cells carrying the monoallelic FANCD2-low deletion also showed delayed tumour growth, with a median time to establishment of 60 days compared with 25 days for wild-type tumours (Figure 5B), paralleling the delayed tumour growth observed upon FBXL12 loss (Figure 2E, 5B). As with FBXL12-KO tumors, excised FANCD2-low tumors showed elevated γH2AX relative to wild-type (Figure S5B), indicating that both FBXL12 loss and FANCD2 insufficiency increase replication-associated DNA damage in vivo.

**Figure 5.**
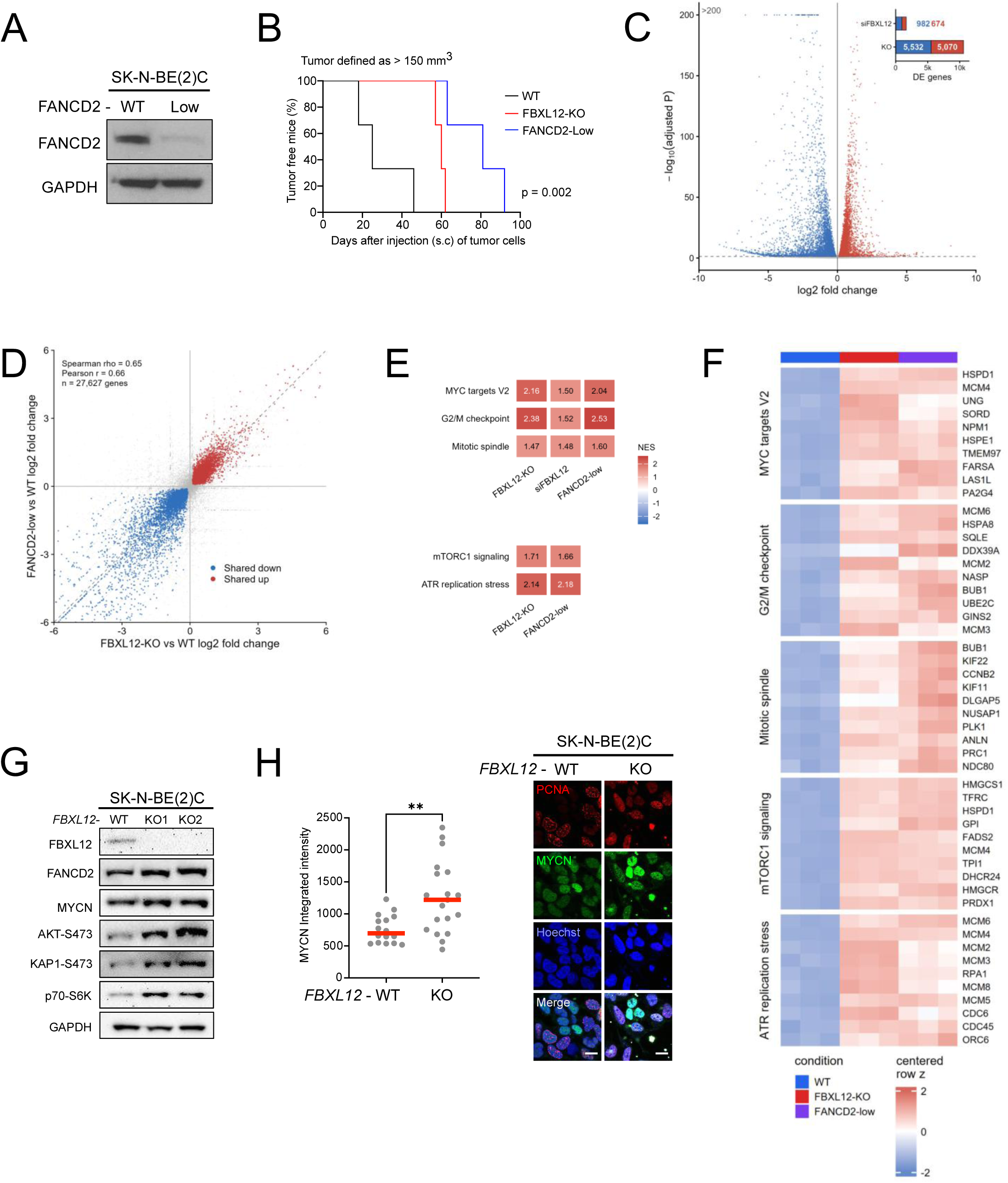
FBXL12 loss and FANCD2 insufficiency converge on a common MYCN-linked stress-adaptation programme. (A) Immunoblots of FANCD2 in total lysates from SK-N-BE(2)C WT and FANCD2-Low clones. (B) Kaplan-Meier curves of the percentage of tumour-free mice following subcutaneous (s.c) injection of SK-N-BE(2)C WT (black, n = 3) FBXL12-KO (red, n = 3) and FANCD2-Low (blue, n = 3) clones. Tumour formation was defined as a tumour volume greater than 150 mm^3^. (C) Volcano plot showing differential gene expression in SK-N-BE(2)C FBXL12-knockout cells compared with wild-type cells. Inset shows the number of significantly upregulated and downregulated genes after acute FBXL12 depletion and chronic FBXL12 knockout. (D) Correlation of differential gene expression changes in FBXL12-knockout versus wild-type cells and FANCD2-low versus wild-type cells. Genes significantly deregulated in both comparisons are highlighted as shared upregulated or shared downregulated genes. (E) Gene set enrichment analysis of selected transcriptional programmes. MYC targets V2, G2/M checkpoint and mitotic spindle signatures are shown across FBXL12-knockout, acute FBXL12-depletion and FANCD2-low conditions; mTORC1 signalling and ATR replication-stress signatures are shown for the chronic FBXL12-knockout and FANCD2-low conditions. Values within tiles indicate normalized enrichment scores (NES), with positive values indicating enrichment in the indicated depleted or knockout condition. Bejamini-Hochberg-adjusted P values were calculated separately within each contrast and gene-set collection and are reported in Table S2. (F) Heatmap of leading-edge genes from the indicated MYC, cell-cycle, mTORC1 and ATR replication-stress gene sets in wild-type, FBXL12-knockout and FANCD2-low cells. Values represent variance-stabilized, row-centred expression. (G) Immunoblot analysis of FBXL12, FANCD2, MYCN, AKT-S473, KAP1-S824 and p70-S6K in SK-N-BE(2)C wild-type cells and FBXL12-knockout clones. (H) Representative immunofluorescence images and quantification of MYCN protein level in PCNA-positive cell in SK-N-BE(2)C WT and FBXL12 KO cells.

To ask whether FBXL12 loss and FANCD2 insufficiency converge at the transcriptional level, we performed RNA sequencing of SK-N-BE(2)C wild-type, FBXL12-knockout and FANCD2-low cells, together with cells subjected to acute siRNA-mediated FBXL12 depletion for 48 hours (Figure S5C). Chronic FBXL12 loss produced substantially more extensive transcriptional remodelling than acute depletion, consistent with progressive adaptation to persistent replication stress, whereas acute FBXL12 depletion showed a more pronounced skew toward gene downregulation (Figure 5C and S5C-D). The FANCD2-low transcriptome showed a striking overlap with FBXL12-knockout cells, supporting convergence on a shared transcriptional state upon chronic disruption of FBXL12-dependent FANCD2 turnover, whether by removing FBXL12 or by limiting the FANCD2 pool itself (Figure 5D and Figure S5E).

Gene set enrichment analysis identified cell-cycle and proliferation-associated programmes, including MYC target genes, G2/M checkpoint and mitotic-spindle signatures, as shared features of FBXL12 loss and FANCD2-low cells. These programmes were already detectable after acute FBXL12 depletion, but were more extensive in chronic FBXL12-knockout and FANCD2-low cells. In contrast, signatures linked to replication-stress adaptation and survival signalling, including Reactome ATR activation in response to replication stress and Hallmark mTORC1 signalling, were most prominently enriched in the chronic knockout and FANCD2-low settings. Thus, acute FBXL12 depletion captures an early cell-cycle response, whereas chronic disruption of FBXL12-dependent FANCD2 turnover selects for a broader adaptive state marked by proliferative, replication-stress and mTOR-associated signalling programmes (Figure 5E, F; Table S2).

Immunoblotting of SK-N-BE(2)C wild-type and two independent FBXL12-knockout clones confirmed the gene set enrichment findings at the protein level, with elevated MYCN, increased p70-S6K, and increased pS33-RPA in FBXL12-knockout cells, consistent with mTOR and ATR pathway activation respectively (Figure 2A and Figure 5G). Since MYCN mRNA is not correspondingly increased in these clones (Figure S2C), this points to a post-transcriptional, protein-level mechanism of MYCN accumulation rather than increased MYCN transcription. Consistent with replication stress and S-phase enrichment, FBXL12-knockout cells showed increased pS473-KAP1, a CHK2-dependent modification induced by replication stress (Lee et al., 2012) and during normal S-phase entry (Chang et al., 2008). Furthermore, immunofluorescence for MYCN in SK-N-BE(2)C wild-type and FBXL12-knockout cells showed higher MYCN intensity in PCNA-positive S-phase cells in FBXL12-knockout relative to wild-type cells (Figure 5H), confirming that the increase in total MYCN protein is reflected specifically within S-phase cells, consistent with the noncanonical fork-associated MYCN function (Solvie et al., 2022).

Together, these data show that disrupting the FBXL12-FANCD2 axis, whether by removing FBXL12 or by limiting FANCD2 itself, converges on a shared transcriptional and signalling state marked by MYC-target, ATR and mTOR pathway activation, accompanied by an increase in S-phase-associated MYCN.

### Disruption of the FBXL12-FANCD2 axis exposes pharmacologically actionable replication-stress and survival-signalling dependencies (Figure 6, Figure S6)

To determine whether mTOR activation contributes to MYCN stabilisation in FBXL12-deficient cells, we treated synchronised SK-N-BE(2)C wild-type and FBXL12-knockout cells released from mitosis into G1 with the mTOR inhibitors everolimus (mTORC1) or AZD2014 (dual mTORC1/2), and assessed MYCN, p70-S6K, and 4E-BP1 by immunoblot (Figure 6A). Both knockout clones showed elevated MYCN, p70-S6K, and 4E-BP1 at baseline. Both inhibitors reduced p70-S6K uniformly, confirming on-target mTOR inhibition, but reduced MYCN more strongly in knockout than in wild-type cells, indicating that knockout cells are more dependent on mTOR-driven MYCN stabilisation. The reduction in 4E-BP1 upon mTOR inhibition was more pronounced in KO2 than KO1 and was not observed in DMSO-treated wild-type cells, indicating clone-to-clone variation in mTOR-dependent 4E-BP1 regulation.

**Figure 6.**
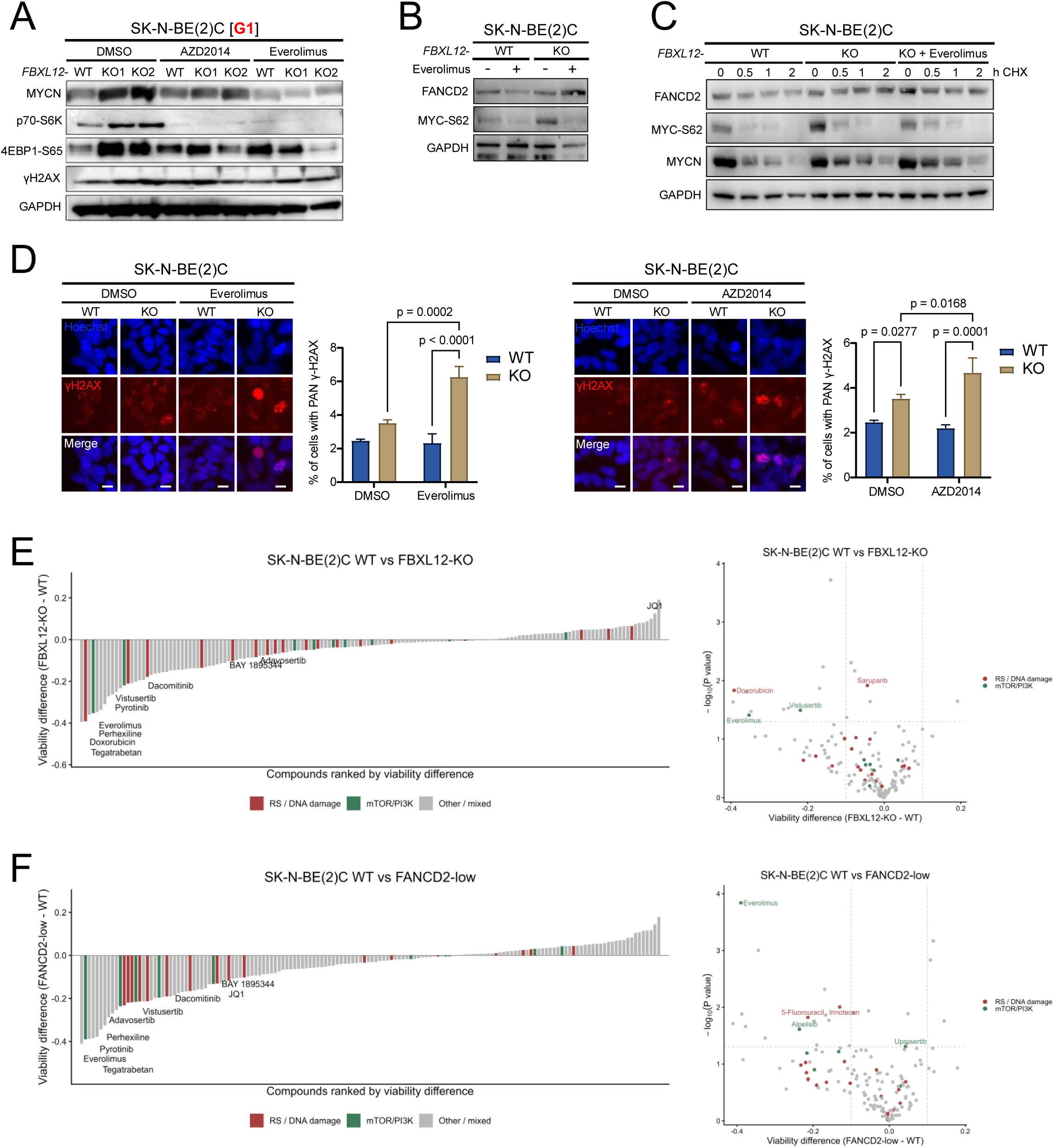
Disruption of the FBXL12-FANCD2 axis exposes pharmacologically actionable replication-stress and survival-signalling dependencies. (A) Immunoblot analysis of MYCN, p70-S6K and 4E-BP1 signalling in SK-N-BE(2)C wild-type and FBXL12-knockout cells treated with the indicated mTOR inhibitors. (B) Immunoblot analysis of FANCD2 and MYCN-S62 phosphorylation in SK-N-BE(2)C wild-type and FBXL12-knockout cells following everolimus treatment. (C) Cycloheximide chase analysis of FANCD2, MYCN-S62 phosphorylation and MYCN protein turnover in SK-N-BE(2)C wild-type and FBXL12-knockout cells, with or without everolimus treatment. (D) Representative γH2AX immunofluorescence images and quantification following everolimus or AZD2014 treatment in SK-N-BE(2)C wild-type and FBXL12-knockout cells. Scale bar, 10 µm. (E) High-content imaging-based compound screen comparing SK-N-BE(2)C wild-type and FBXL12-knockout cells. Left, compounds ranked by viability difference between FBXL12-knockout and wild-type cells, with negative values indicating lower viability in FBXL12-knockout cells. Right, volcano plot showing viability difference against −log10(P value). Compounds targeting replication-stress/DNA-damage pathways and mTOR/PI3K signalling are highlighted in red and green, respectively. (F) Corresponding high-content imaging-based compound screen analysis comparing SK-N-BE(2)C wild-type and FANCD2-low cells. Left, compounds ranked by viability difference between FANCD2-low and wild-type cells, with negative values indicating lower viability in FANCD2-low cells. Right, volcano plot showing viability difference against −log10(P value), with pathway categories coloured as in (E). Dashed lines indicate P=0.05 and viability differences of ±0.10.

To determine whether this mTOR-dependent stabilisation acts through the canonical MYCN degradation pathway, we next examined the stabilising Ser62 phosphorylation mark, which precedes GSK3β/FBXW7-dependent degradation at Thr58 (Farrell and Sears, 2014). Baseline pS62-MYCN was elevated in FBXL12-knockout relative to wild-type cells, consistent with MYCN stabilisation in knockout cells in line with their elevated mTOR pathway activity (Figure 6B). Everolimus reduced pS62-MYCN to a similar extent in wild-type and FBXL12-knockout cells, confirming on-target suppression of active MYCN in both genotypes, but FANCD2 levels were unaffected, remaining elevated in knockout relative to wild-type (Figure 6B).

To determine whether FANCD2 turnover depends on MYCN or mTOR activity, we performed cycloheximide (CHX) chase in wild-type, FBXL12-knockout, and everolimus-treated FBXL12-knockout cells. FANCD2 showed partial decay within 2 hours in wild-type cells but remained stable in FBXL12-knockout cells regardless of mTOR inhibition (Figure 6C) indicating that FANCD2 stabilization is independent of mTOR activity. MYCN turnover was only slightly delayed in knockout cells but remained substantially degraded within the same timeframe, and pS62-MYCN decayed similarly across all conditions (Figure 6C). This dissociation between FANCD2 stability and ongoing MYCN/pS62-MYCN turnover suggests that FANCD2 degradation does not simply track with MYCN abundance. Everolimus and AZD2014 nonetheless induced significantly greater γH2AX in FBXL12-knockout than wild-type cells (Figure 6D), indicating that mTOR pathway activity contributes to replication-stress tolerance in knockout cells, such that its inhibition unmasks additional DNA damage independently of any effect on FANCD2 stability.

We next asked whether these pathway dependencies extended to differential drug sensitivity more broadly. A high-content imaging drug-sensitivity screen of 150 FDA-approved and investigational compounds in SK-N-BE(2)C wild-type, FBXL12-knockout, and FANCD2-low cells, scored by cell viability, identified genotype-dependent shifts across a subset of pathway classes (Figure 6E–F and table S3). The most consistent, statistically significant hits across both FBXL12-knockout and FANCD2-low comparisons targeted mTOR (everolimus, vistusertib), consistent with mTOR-dependent stabilisation of Ser62-phosphorylated MYCN (Vaughan et al., 2016), ERBB-family receptor tyrosine kinases (pyrotinib, dacomitinib, perhexiline), which converge on the ERK-Ser62 axis, and Wnt/β-catenin signalling (tegatrabetan). A smaller set of TEAD/Hippo pathway inhibitors also showed reproducible genotype-selective sensitivity, in line with a TEAD4–MYCN feedback loop described in high-risk neuroblastoma (Rajbhandari et al., 2018). In line with the established function of FANCD2, FANCD2-low cells were more sensitive to BET/MYC bromodomain inhibition (JQ1) (Figure 6E–F and table S3). ATR inhibitors did not reach significance in either FBXL12-KO or FANCD2-low cells, although ATR pathway activation was evident from gene set enrichment analysis and immunoblot (Figure 5E–G). We therefore performed a secondary, focused high-content imaging screen and assessed EdU incorporation and γH2AX induction in FBXL12-knockout versus wild-type cells for a curated panel of compounds, selected to include candidates targeting DNA damage response and replication-stress pathways implicated by RNA-seq and immunoblot (Figure 5E–G). Matching each compound to the dose producing the largest knockout-directional shift, the CHK1 inhibitor rabusertib and the topoisomerase I inhibitor LMP744 produced the most pronounced knockout-selective effects, combining increased γH2AX-positive foci and reduced S-phase occupancy, a signature consistent with checkpoint-driven exit from S phase following unresolved damage (Figure S6A and table S3). The ATR inhibitor BAY 1895344, the DNA-damaging agents cisplatin and decitabine, and the PARP inhibitor talazoparib showed a similar, coherent pattern. A small number of compounds, including the BCL-2/BCL-XL inhibitor navitoclax, showed no knockout-selective damage induction, arguing that the knockout-directional effects were not a general consequence of reduced proliferation in FBXL12-deficient cells. Together, the two independent screens converge on mTOR signalling and replication-stress/DNA-damage response pathways as the most reproducible drug-sensitivity signature of FBXL12 deficiency, without establishing single-agent efficacy.

As an orthogonal cell-line analysis, we next asked whether endogenous FBXL12 expression was associated with mTOR-inhibitor response in DepMap drug-sensitivity data (Ghandi et al., 2019, Corsello et al., 2020). Across 13 neuroblastoma cell lines, FBXL12 expression positively correlated with mTOR-inhibitor AUC values, indicating that cell lines with lower FBXL12 expression tended to show greater sensitivity to mTOR-pathway inhibition. This association was observed for two mTOR inhibitors, reaching statistical significance for one compound (Figure S6B). Although limited by the number of neuroblastoma models available, these data are consistent with enhanced mTOR-inhibitor sensitivity in FBXL12-low or FBXL12-disrupted cells.

Finally, to relate these experimental models back to the clinical association between high FBXL12 expression and poor clinical outcome, we asked whether endogenous FBXL12-expressing neuroblastoma cells approximate the FBXL12-high state observed in patient tumours. To reduce scale differences between datasets, we recalculated gene-level RPM values from the SK-N-BE(2)C RNA-seq count matrix and compared FBXL12 log2RPM values with the processed SEQC neuroblastoma cohort. Although this does not remove cross-study batch, processing, annotation, or tumour-versus-cell-line differences, FBXL12 expression in SK-N-BE(2)C wild-type and siSCR control cells fell within the upper range of SEQC tumours, whereas siFBXL12 depletion reduced FBXL12 expression below the patient median (Figure S6C). In this context, FBXL12 knockout and siRNA depletion provide functional perturbations of an endogenous FBXL12-high/intact state.

Together, these data indicate that disrupting the FBXL12-FANCD2 axis exposes candidate pharmacological dependencies converging on mTOR, ATR, and MYC-directed signalling, nominating pathways that may become therapeutically exploitable when this genome-maintenance axis is disrupted in FBXL12-high tumours.

## DISCUSSION

Our results identify FBXL12-mediated control of FANCD2 turnover as a replication stress tolerance mechanism that is co-opted by MYCN in neuroblastoma. As previously shown in cyclin E-driven breast cancer (Brunner et al., 2023), timely removal of FANCD2 from chromatin is required to prevent a fork-protective factor from becoming a source of replication-associated damage. Here, we extend this principle to MYCN-driven neuroblastoma and beyond S phase, showing that defective FANCD2 turnover also impairs MiDAS and is associated with the transmission of unresolved DNA lesions into daughter-cell G1.

Together, these findings support a model in which MYCN regulates the balance of chromatin-bound FANCD2 through FBXL12, and in which disruption of this balance compromises the processing of replication intermediates across both S phase and mitosis (Figure 7). In FBXL12-intact, MYCN-amplified cells, MYCN-driven replication stress is met by FBXL12-mediated turnover of chromatin-bound FANCD2, which is licensed by CHK1-dependent phosphorylation of FANCD2 (Brunner et al., 2023). MYCN itself antagonises this degradation, so that the two activities together maintain a balanced, chromatin-associated FANCD2 pool. This balance supports fork protection and restart in S phase and is compatible with FANCD2-dependent MiDAS at under-replicated loci during mitosis (Figure 7). In FBXL12-deficient, MYCN-amplified cells, loss of this degradation step allows FANCD2 to accumulate and persist on chromatin. Rather than further enhancing fork protection, this excess is associated with impaired fork progression, ATR-dependent replication-stress signalling, and reduced MiDAS. Our epistasis data (Figure 3D-G) best support this model, though endpoint-based assays cannot exclude a role for MYCN in initial FANCD2 recruitment. Because this model identifies FBXL12, the FBXL12-MYCN interaction, and FANCD2 itself as the nodes that set this balance, each represents a rationally distinct point at which the axis could be pharmacologically disrupted (Figure 7).

**Figure 7.**
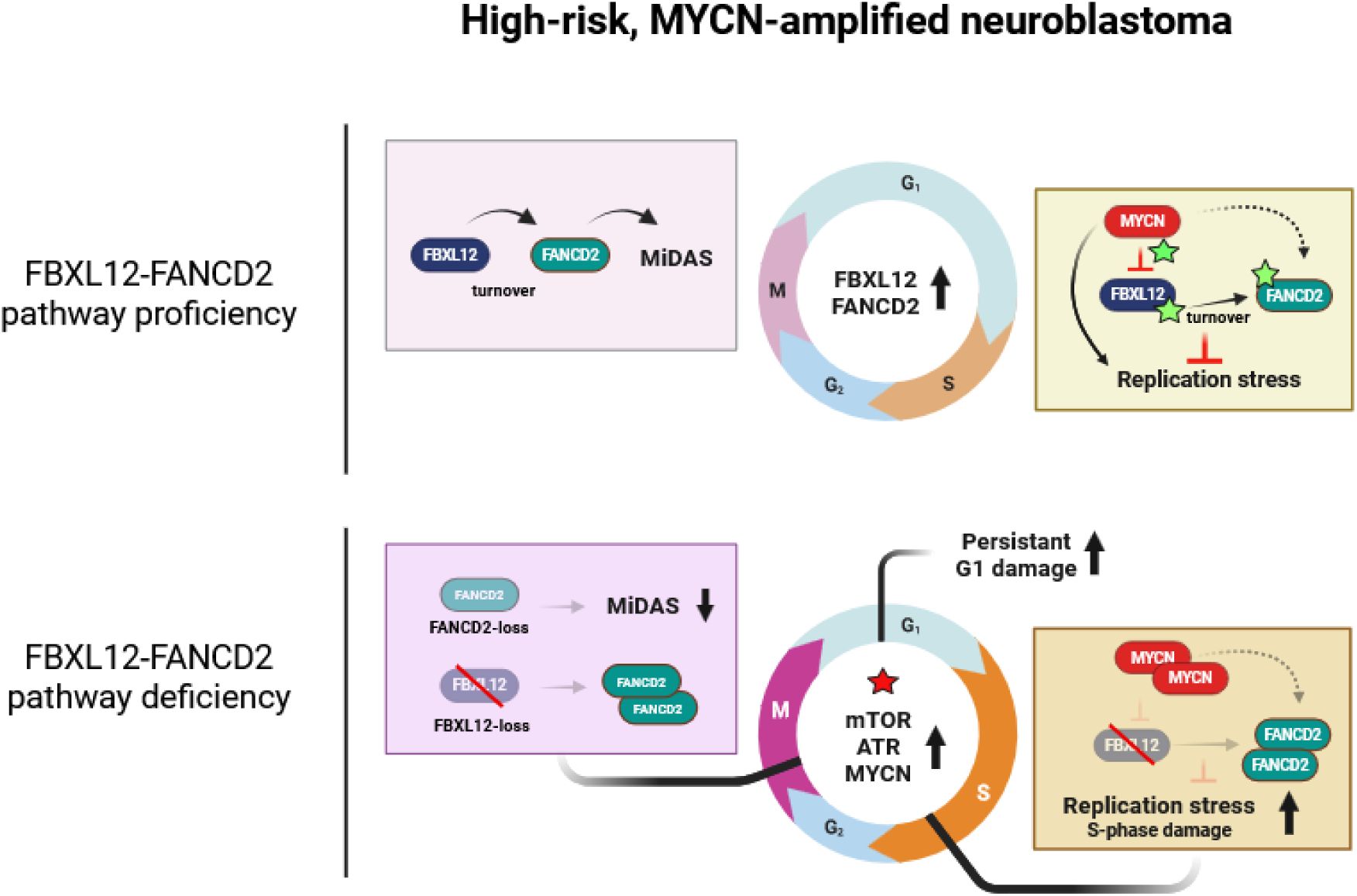
Schematic of the MYCN-FBXL12-FANCD2 axis and its disruption in FBXL12-deficient, MYCN-amplified neuroblastoma. In pathway-proficient cells (top), FBXL12-mediated FANCD2 turnover supports fork restart and mitotic DNA synthesis (MiDAS). MYCN acts upstream of FBXL12 maintaining a balanced, transiently chromatin-bound FANCD2 pool. Stars mark the three actionable nodes setting this balance: MYCN-FBXL12 interaction, FBXL12 activity, and FANCD2 itself. In pathway-deficient cells (bottom), FBXL12 loss uncouples this balance and FANCD2 instead accumulates and persists on chromatin. Unlike simple FANCD2 loss, which abolishes MiDAS (left inset, top), FBXL12 loss converts a transient, productive FANCD2-chromatin association into a persistent, non-productive one (left inset, bottom), reducing MiDAS while elevating S-phase stress and damage that is transmitted through mitosis into G1. Compensatory mTOR, ATR and MYCN signalling (starred hub) nominates further pharmacological vulnerabilities exposed by this disruption.

While FANCD2 loss itself abolishes MiDAS entirely (Xu et al., 2021), FANCD2 foci remain detectable in prometaphase despite reduced mitotic DNA synthesis in FBXL12-deficient cells. Thus, FBXL12 loss does not simply phenocopy FANCD2 loss but instead converts a normally transient, productive chromatin association into a persistent, non-productive one. We therefore propose that persistent FANCD2 becomes non-productive when it is not removed at the appropriate stage of replication or mitosis. Such persistence could prevent the remodelling of replication intermediates or restrict access by downstream MiDAS factors, including the SLX4-MUS81–EME1, RAD52, and POLD3 machinery (Minocherhomji et al., 2015, Bhowmick et al., 2016).

FANCD2 depletion rescues both S-phase and G1 damage in FBXL12-deficient cells, identifying dysregulated FANCD2 as an upstream determinant of both phenotypes. MYCN depletion, by contrast, rescues S-phase damage but not the G1 lesions detected under the conditions examined. These observations demonstrate that the two phenotypes differ in their response to MYCN depletion and suggest that MYCN is mechanistically less relevant to the generation of G1 damage in FBXL12-deficient cells. However, G1 lesions may originate from replication intermediates formed during an earlier period of MYCN-driven stress, before MYCN depletion can reverse or prevent their transmission through mitosis. Thus, although our epistasis experiments support differential dependence of the measured S-phase and G1 phenotypes on ongoing MYCN activity, further temporal experiments will be needed to determine when MYCN acts in relation to the formation and mitotic processing of these lesions.

These data extend our previous description of FBXL12-mediated FANCD2 degradation as a replication-recovery mechanism in cyclin E-driven breast cancer (Brunner et al., 2023) to an oncogene-driven paediatric cancer context. Here, rather than FANCD2 turnover being determined solely by FBXL12 activity, it is also controlled by the oncogenic driver that generates the replication stress in the first place. This places MYCN in a dual role at the replication fork. MYCN generates replication stress through its transcriptional and replicative activity while simultaneously acting, at least in part through its interaction with FBXL12, to preserve a chromatin-bound pool of FANCD2 sufficient to tolerate that stress (Figure 7).

This mechanism is conceptually related to, but mechanistically distinct from, the recently described noncanonical capacity of MYC to multimerise at stalled forks and physically shield them from transcribing RNA polymerase, an effect that similarly enhance FANCD2 chromatin association (Solvie et al., 2022). In the same study, depletion of CHK1 or FANCI, but not of core double-strand-break repair factors including BRCA1, BRCA2, ATM, or MRE11, reduced MYC multimer formation, implicating the same CHK1– FANCD2/FANCI axis identified here as a determinant of MYC multimerisation at stalled forks. Our data suggest an additional, degradation-based mechanism that converges on the same outcome, chromatin-retained FANCD2, and raise the question of whether MYC/MYCN multimerisation and FBXL12 antagonism reflect a shared CHK1-licensed fork-protective programme or parallel, additive mechanisms of replication-stress tolerance. Resolving this will require testing whether MYCN multimerisation itself, rather than MYCN abundance alone, is required to protect FANCD2 from FBXL12-mediated degradation. This picture is consistent with independent reports of stress-responsive, non-transcriptional MYC-family engagement with the DNA damage response. MYCN adopts a distinct physical state under S-phase and transcriptional stress (Papadopoulos et al., 2022, Papadopoulos et al., 2024), interacting with nuclear exosome-targeting complexes that resolve transcription-replication collisions and eliminate genotoxic RNA structures such as R-loops (Papadopoulos et al., 2022, Papadopoulos et al., 2024). Our FBXL12 interactome did not recover nuclear exosome or R-loop-associated factors, suggesting this represents a mechanistically distinct MYCN pool from the FBXL12-FANCD2 axis described here, consistent with reports that MYC-family proteins partition into functionally separable pools with distinct stabilities (Lourenco et al., 2021). MYC’s association with FANCD2 and DNA damage sites also intensifies under replication stress in a manner dependent on Ser 62 phosphorylation (Cohn et al., 2026), the same phosphodegron-priming mark that is elevated in our FBXL12-knockout cells (Figure 6B). This raises the possibility that the elevated pS62-MYCN pool in FBXL12-deficient cells contributes to replication-stress tolerance through a repair-recruitment function in addition to its role in shielding FANCD2 from degradation, and may explain why mTOR inhibition, which reduces pS62-MYCN, selectively increases DNA damage in knockout cells (Figure 6B and 6D).

The clinical findings suggest that this pathway may represent an adaptation of aggressive neuroblastoma rather than a conventional tumour-suppressive relationship. FBXL12 expression is elevated in high-risk and MYCN-amplified neuroblastoma and independently predicts inferior survival across patient cohorts (Figure 1), whereas experimental FBXL12 loss compromises the fitness of MYCN-amplified cells. This is consistent with MYCN-amplified tumours maintaining high FBXL12 expression because regulated FANCD2 turnover is required to tolerate their elevated replication-stress burden. High FBXL12 marks and supports an aggressive cellular state, while acute FBXL12 loss exposes a dependency of that state. Our model identifies several distinct points at which the MYCN-FBXL12-FANCD2 balance could be pharmacologically disrupted (Figure 7). Our data show that FBXL12 loss alone compromises the fitness of MYCN-amplified cells both in vitro and in vivo, providing direct rationale for its catalytic inhibition. A second, potentially more selective node is the FBXL12-MYCN interaction itself. A related precedent exists for MYCN where AURKA protects MYCN from FBXW7-mediated degradation through a direct, kinase-independent interaction, and disrupting this interaction pharmacologically promotes MYCN degradation in neuroblastoma (Otto et al., 2009, Richards et al., 2016). By analogy, agents that disrupt the MYCN-FBXL12 interaction could restore FANCD2 degradation and sensitise MYCN-amplified cells to replication stress without broadly inhibiting FBXL12 elsewhere. A third node is FANCD2 itself. Reducing the chromatin-bound FANCD2 pool, and not only preventing its removal, similarly impairs fitness, nominating FANCD2 degradation as a potential route to disrupting this balance. Disrupting the FBXL12-MYCN-FANCD2 axis also exposes additional vulnerabilities, including mTOR, ERBB and ATR/CHK1 activities, consistent with the elevated replication stress and mitotic DNA synthesis defect that follow FBXL12 loss (Figure 6). Together, FBXL12 catalytic activity, the FBXL12-MYCN interaction, FANCD2 stability, and the compensatory ATR/mTOR/MYC signalling exposed by their disruption, offer complementary and potentially combinable strategies for targeting this axis (Figure 7), rather than pointing to a single validated drug target.

Together, our findings establish MYCN as a regulator of FBXL12-mediated FANCD2 turnover and extend the consequences of defective turnover from replication recovery in S phase to MiDAS and inherited G1 damage. They support a model in which MYCN both generates replication stress and preserves the FANCD2 pool required to manage it, while FBXL12 prevents that protective pool from becoming persistently and deleteriously associated with chromatin. Disrupting this balance compromises replication-fork progression, impairs the mitotic processing of under-replicated DNA, and is associated with the transmission of DNA lesions into daughter cells. More broadly, these findings show how an established mechanism of replication recovery can be co-opted by an oncogenic driver and expose regulated FANCD2 turnover as a potential vulnerability of MYCN-amplified tumours.

## Supporting information

Table S1 MS results fig 3A

Table S2 GSEA

Table S3 drug screens

Table S4 antibodies

Table S5 oligos

Table S6 xenograft data

## ACKNOWLEDGEMENTS

Work in the O.S. laboratory is supported by the Swedish Cancer Society, the Swedish Childhood Cancer Fund, the Swedish Research Council, Karolinska Institutet, Radiumhemmets Research Foundation, and Worldwide Cancer Research (grant reference number 26-0033). A.B.’s salary is supported by the Worldwide Cancer Research grant. Research led by M.W. and J.I.J. is supported by the Swedish Cancer Society, the Swedish Childhood Cancer Fund and Radiumhemmets Research Foundation. L.M.O. receives funding from the Swedish Cancer Society (22 2000 Pj, 25 4349 Pj), the Swedish Research Council (2024-03294), Radiumhemmets Research Foundation (244203) and Karolinska Institutet (2023-01372). Work in the M.P. laboratory is supported by Karolinska Institutet, the Swedish Research Council and the Swedish Cancer Society. We also acknowledge the AKM Animal Core Facility for their support with animal experiments and Anna Malmerfelt at Histology Services, Theme Cancer, for technical assistance. The authors thank Drs. Fredrik Swartling and Margareta Wilhelm for kindly providing MYCN plasmids and Dr. Laura Baranello for critical reading.

## Material and methods

### Mammalian cell culture

HEK293, HEK293T, SHEP and IMR32 cells were grown in Dulbecco’s modified Eagle’s medium (DMEM) supplemented with 10% FBS and 2Mm L-glutamine. SK-N-BE(2)-C and Kelly cells were grown in RPMI-1640 supplemented with 10% FBS and 2mM L-glutamine. Cell lines were authenticated by short tandem repeat analysis, and all cell lines were mycoplasma-free validated by PCR.

### Antibodies and biochemical analysis

NP-40 lysis buffer (50 mM Tris-HCl [pH 8.0], 150 mM NaCl, 1% NP-40) supplemented with both protease (Complete mini) and phosphatase (PhosSTOP) was used to obtain whole cell lysates from cells. For immunoprecipitation (IP), lysates were incubated with primary antibodies for 16 h at 4◦C, followed by 1 h incubation with Dynabeads coupled with protein G and three times wash by lysis buffer. Proteins were separated by SDS-PAGE on 4-12% bis-acrylamide gels and transferred on to PVDF membranes. The immunoblots were detected by using primary antibodies and subsequent chemiluminescence isotype-specific secondary antibodies coupled to horseradish peroxidase. Antibodies used in the study were listed in Table S4.

### Gene silencing and transfections

Transfection was performed according to the manufacturer’s instruction of the reagent used in the study. siRNA transfection was performed by using Lipofectamine RNAiMAX or HiPerfect, and plasmid transfection was performed by using Lipofectamine 3000. Cells were transfected 48 to 72 h prior to biochemical or functional analysis.

### CRISPR genome editing

Two sgRNAs (Table S5) were designed to target *FBXL12* or *FANCD2* genes and cloned each of them into the *Bbs*I site of pSp-Cas9-GFP vector. SK-N-BE(2)-C and Kelly cells were transfected with 1 µg/ml of each pSp-Cas9-GFP-sgRNA vector or mock transfected with empty pSp-Cas9-GFP vector. A total of 48 h post transfection, cells were resuspended and seeded into 96-well plate at the density of 1-5 cells/well to grow single cell clones. Genomic DNA of individual clones was screened by PCR (Table S5) and the deletion was validated by Sanger sequencing.

### Flow cytometry

Kelly and SK-N-S/BE(2)-C cells were pulse-labelled with EdU for 30 min, washed, and released into EdU-free medium. Cells were collected at the indicated time points following release and processed for flow cytometry. EdU incorporation was detected together with Hoechst staining to determine DNA content. Cell-cycle progression of the EdU-labelled population was assessed based on combined EdU incorporation and DNA content. EdU-positive cells with DNA content close to 2N were classified as early S phase, EdU-positive cells approaching 4N DNA content were classified as late S phase. EdU-negative cells with 2N and 4N DNA content were classified as G1 and G2/M, respectively. The percentage of cells within each gate was quantified at each time point. For analysis of cell-cycle progression, the late S and G2/M populations were plotted over time. Area under the curve (AUC) was calculated using the trapezoidal method from the original measured time points within the indicated interval.

### High content imaging-based drug screening

High-content imaging screens were performed in SK-N-BE(2)C wildtype and FBXL12-knockout cells. In the primary screen, 150 FDA-approved and investigational compounds were tested at four concentrations. A secondary screen evaluated a focused panel of 40 compounds at four concentrations. Following compound treatment, cells were labelled with EdU, fixed and permeabilized. EdU incorporation was detected by click chemistry, and γH2AX was detected by immunofluorescence. Nuclei were counterstained with DAPI.

Images were acquired by automated high-content microscopy. DAPI staining was used for nuclear segmentation and determination of relative cell viability, while EdU incorporation was used to quantify proliferating cells and cell-cycle distribution. DNA-damage responses were assessed from γH2AX staining, including nuclear γH2AX intensity and γH2AX-positive foci. Genotype-dependent responses were calculated as the difference between FBXL12-knockout and wildtype cells.

### Colony formation assay

SK-N-BE(2)-C cells were seeded into 6-well plates at the density of 1000 cells/well. The colony formation was determined after 2 weeks post seeding. Colonies were recorded by a digital camera and counted by Fuji ImageJ.

### Mass spectrometry

Bead-bound proteins were reduced, alkylated, and digested overnight with sequencing-grade modified trypsin. Peptides were purified using a modified SP3 protocol (Moggridge et al., 2018) and analyzed by nano-flow liquid chromatography coupled to a timsTOF HT mass spectrometer using DIA-PASEF acquisition.

Nine samples were analyzed, comprising three replicate control immunoprecipitations under MG132-treated conditions, three replicate FBXL12 immunoprecipitations under untreated conditions, and three replicate FBXL12 immunoprecipitations under MG132-treated conditions.

Proteins were identified and quantified at the protein-group level. Candidate FBXL12 interactors were evaluated by comparing replicate-averaged MS2 quantities between FBXL12 and control immunoprecipitates. Enrichment was expressed as fold change and log₂ fold change together with calculated p-values. The effect of MG132 treatment was assessed by comparing treated and untreated FBXL12 immunoprecipitates.

### Sub-cellular fractionation

Fractionation of chromatin-associated proteins was performed as previously described (Pena-Diaz et al., 2012). Cell suspension was incubated with pre-extraction buffer for 5 min and spun down. The supernatant part was kept as soluble fraction. The cell pellet was resolubilised in NP-40 buffer, followed by sonication and spin down. The supernatant was collected as chromatin-enriched fraction and subjected to analysis described above.

### Analysis of proteins on nascent DNA

Analysis of proteins associated with nascent DNA was carried out using a modified accelerated native isolation of proteins on nascent DNA (aniPOND) protocol adapted from (Leung et al., 2013). Briefly, cells, approximately 4 × 10⁷ cells per condition, were pulse-labeled with 10 μM EdU for 0.5 h. Nuclei were then prepared using nuclei extraction buffer containing 20 mM HEPES, pH 7.2, 50 mM NaCl, 3 mM MgCl₂, 300 mM sucrose, and 0.5% IGEPAL CA-630. Biotin-Azide click chemistry was performed on isolated nuclei with rotation for 1 h at 4 °C.

After the click reaction, nuclei were lysed in buffer containing 25 mM NaCl, 2 mM EDTA, 50 mM Tris-HCl, pH 8.0, 1% IGEPAL CA-630, and protease inhibitors. Chromatin was briefly sheared by sonication for 10 s at maximum power using a Diagenode Bioruptor, followed by centrifugation and removal of the supernatant. This extraction step was repeated once to deplete soluble and non-chromatin-bound proteins. The resulting chromatin pellet was resuspended in lysis buffer and subjected to 12 cycles of sonication, each for 10 s at maximum power, to release chromatin-associated proteins. The salt concentration of the final chromatin lysate was adjusted to physiological levels by adding lysis buffer containing 150 mM NaCl. Streptavidin pull-down was performed using 50 μl of streptavidin beads per sample. After three washes, bead-bound proteins were eluted by boiling in 1× SDS sample loading buffer and analyzed by electrophoresis.

### Immunofluorescence staining and microscopy

Cells were seeded on iBidi Chambered coverslip (iBidi µ-Slide 18 Well, 81816) or Corning Falcon clear microplate (high content imaging) for the indicated experiments. Then cells were subjected to fixation with 4% paraformaldehyde for 10 min and permeabization/blocking with 0.1% Triton-X 100, 5% bovine serum albumin (BSA) in TBS for 1 h at room temperature. Subsequently, cells were incubated with primary antibodies of the interested proteins in 5% BSA TBS for 16 h at 4℃ and then washed three times with TBS-T, followed by incubation with appropriate isotype-specific secondary antibodies coupled different fluorescence and Hoechst 33342 stain. After washing three times with TBS-T, cells were mounted by ProLong Diamond antifade mounting solution (coverslip) or PBS (Corning Falcon clear microplate). Images were taken using EVOS™ M7000 Imaging System and analysed by using CellProfiler 4.2.8 and Graphpad Prism 11.

### Mitotic DNA synthesis (MiDAS) assay

Cells were seeded in 10-cm dishes 24 h before synchronization with 2 mM thymidine for 16 h. The cells were then washed three times with prewarmed DPBS and incubated with RO-3306 for an additional 16 h. During the final hour of RO-3306 treatment, 10 µM mirin was added as a positive control. The synchronized cells were gently washed three times with prewarmed DPBS and incubated for 30 min in prewarmed medium containing 20 µM EdU and 10 µM mirin to allow progression into prometaphase. Prometaphase cells were collected by mitotic shake-off and seeded onto chambered coverslips (µ-Slide 18 Well, 81816; ibidi). After allowing the cells to settle for 5 min, they were fixed with 4% paraformaldehyde for 10 min. The cells were then permeabilized and blocked for 10 min at room temperature in TBS containing 0.1% Triton X-100 and 5% bovine serum albumin. EdU labeling was detected using the Click-iT Plus EdU Alexa Fluor 488 Kit according to the manufacturer’s instructions. Following the click reaction, the cells were immunostained for the proteins of interest as described above. Images were acquired using a Zeiss LSM 800-Airy microscope and analyzed using CellProfiler version 4.2.8 and GraphPad Prism version 11.

### RNA sequencing and differential expression analysis

Total RNA was extracted from SK-N-BE(2)C neuroblastoma cells using the QIAGEN RNeasy Plus kit. RNA integrity was confirmed prior to sequencing, with all samples showing RIN values between 9.4 and 10. Bulk mRNA sequencing was performed on 18 samples, with three biological replicates per condition: siSCR, siFBXL12, siFANCD2, wild-type, FBXL12-knockout, and FANCD2-low cells. Libraries were prepared using the Illumina TruSeq Stranded mRNA kit with poly(A) selection and sequenced on a NovaSeq X Plus using paired-end 2 × 150 bp sequencing, targeting approximately 30 million reads per sample.

Reads were aligned and quantified against the human GRCh38 reference genome, and downstream analyses were performed from a STAR/Salmon gene-level count matrix. Salmon gene counts were rounded to integers and analysed in R using DESeq2 v1.46.0. Genes with Benjamini-Hochberg adjusted P value < 0.05 were considered differentially expressed.

Gene set enrichment analysis was performed using clusterProfiler v4.14.6 and msigdbr v25.1.1. Genes were ranked by the DESeq2 Wald statistic after mapping Ensembl IDs to HGNC symbols. GSEA was performed using MSigDB Hallmark gene sets and, where indicated, curated C2 gene sets. Leading edge genes were visualized using DESeq2 variance-stabilized expression values centred to the mean expression in wild-type samples.

### Quantitative real-time (qRT) PCR

For qRT-PCR analysis, total RNA was isolated using the RNeasy Kit (Qiagen). Genomic DNA contamination was removed by treating the RNA with DNase I (Qiagen) in accordance with the manufacturer’s instructions. Subsequently, 50 ng of purified RNA was reverse-transcribed into cDNA using the SuperScript VILO cDNA Synthesis Kit (Invitrogen, Life Technologies). Quantitative PCR was performed using SYBR Green chemistry. The primer sequences are provided in Table S5. Relative gene expression was calculated using the ΔΔCt method.

### STRING network analysis of enriched proteins

Proteins enriched in the immunoprecipitation mass spectrometry were analyzed using the STRING functional protein association database. Gene symbols were first mapped to human STRING identifiers using Homo sapiens as the reference organism (NCBI taxonomy ID: 9606).

Protein-protein association data were retrieved from STRING v12.0. Interactions among proteins in the submitted list were retained when the STRING combined interaction score was ≥0.4, corresponding to medium-confidence associations. Network clustering was performed on the retained STRING interaction network. Proteins were grouped according to network connectivity and functional enrichment of the resulting modules. For each cluster, enriched biological annotations were examined using STRING functional enrichment results, including Gene Ontology biological process terms, pathway annotations, protein domains, and curated functional categories. Enrichment terms were filtered to remove broad, non-informative categories.

### Bioinformatics and survival analysis of public neuroblastoma cohorts

Expression and clinical data for neuroblastoma cohorts were obtained from the R2 Genomics Analysis and Visualization Platform, including the Cangelosi multi-cohort dataset, Kocak cohort, and SEQC cohort (Cangelosi et al, 2020; Kocak et al., 2013; Zhang et al., 2015). In the Cangelosi dataset, high-risk status was reconstructed because no explicit annotation was available. Tumours were classified as high risk if they were MYCN-amplified or INSS stage 4 in patients aged ≥18 months. Tumours meeting neither criterion were classified as non-high risk.

Replication-stress activity was assessed using the published Takahashi replication-stress gene-expression signature (Takahashi et al., 2022). Signature scores were calculated from standardized expression values and compared between risk groups and MYCN groups.

KEGG pathway enrichment analysis was performed in the R2 platform using the Cangelosi neuroblastoma dataset after stratifying tumours into reconstructed high-risk and non-high risk groups. Pathways enriched among genes upregulated in high-risk tumours were plotted according to enrichment fraction and Benjamini-Hochberg-adjusted *p* value.

For survival analyses, FBXL12 and FANCD2 expression values were standardized within each cohort. When multiple probes mapped to one gene, probe z-scores were averaged. Kaplan-Meier curves were generated after median dichotomization of expression, with low expression defined as ≤ median and high expression as > median. Survival differences were assessed using two-sided log-rank tests.

Multivariable Cox proportional hazards models were fitted in the Kocak and SEQC cohorts to test whether gene expression was independently associated with outcome. Expression was analysed continuously as a within-cohort z-score. Models were adjusted for age >18 months, MYCN amplification, and stage 4 disease. Stage 4S was included in the non-stage-4 reference group. Hazard ratios therefore represent the effect per 1 SD higher gene expression after adjustment for these clinical risk factors. Proportional-hazards assumptions were assessed using Schoenfeld residuals.

### DepMap Pharmacogenomic Analysis

Neuroblastoma cell-line models were identified from DepMap model annotations. FBXL12 mRNA expression was extracted from the DepMap RNA expression matrix and linked to PRISM drug-response data using stable DepMap model identifiers. Drug sensitivity was assessed using PRISM area under the dose-response curve (AUC). Associations between FBXL12 expression and drug response were evaluated for each compound using Spearman rank correlation. P values were adjusted for multiple testing using the Benjamini-Hochberg method. For group-based comparison, cell lines were divided into FBXL12-high and FBXL12-low groups according to median FBXL12 expression, and AUC values were compared using a two-sided Mann-Whitney U test. Effect direction was interpreted considering that lower AUC represents higher drug sensitivity.

### Subcutaneous xenograft experiments

Five million SK-N-BE(2)C wild-type, FBXL12-knockout or FANCD2-low cells were resuspended in serum-and antibiotic-free RPMI-1640 medium and injected subcutaneously into NMRI nu/nu mice (5–6 weeks old; Taconic) under isoflurane anaesthesia (2-4%), in a final injection volume of 100 µL. The wild-type and FBXL12-knockout groups were studied across two independent experiments comprising a pilot cohort (n = 3 per group) and a prospectively planned follow-up cohort (n = 6 per group). Data from the two experiments were pooled as prespecified, giving a combined sample size of nine mice per genotype. The pilot experiment additionally included three mice injected with FANCD2-low cells. Individual mouse data are provided in Table S6.

Mice were monitored for tumour development and tumour size was measured three times per week using a caliper. Tumour volumes were calculated as width² × length × 0.44. Tumour establishment was defined prospectively as attainment of a tumour volume of 150 mm³. Time from injection to tumour establishment was compared using the log-rank test. Animals whose tumours did not reach the specified volume during follow-up were censored at the end of the observation period. To account for potential variation between the pilot and follow-up experiments, pooled wild-type and FBXL12-knockout data were additionally analysed using a Cox proportional hazards model stratified by experiment. Robustness was assessed by repeating the time-to-event analysis using additional tumour-volume thresholds.

Mice were euthanised when tumour volume exceeded 1500 mm³ or 20 mm, after which tumours were excised, weighed, and processed for downstream analyses, including immunoblot analysis of FBXL12 and γH2AX. Animal health, including body weight, was assessed regularly. Mice were housed in standard cages in a temperature-and humidity-controlled room under a 12h light/12h dark cycle, with ad libitum access to sterile food and water. All procedures were approved by the Stockholm Ethics Committee for Animal Research (permit no. 4356-2024), under the Swedish Board of Agriculture, and were conducted in accordance with Swedish national and EU regulations.

### Statistical analysis

Data were analyzed using GraphPad Prism 11. Unless otherwise stated in the figures or figure legends, results are representative of at least three independent biological replicates and are presented as the mean ± SD. Comparisons between two groups were performed using an unpaired Student’s t-test unless specified otherwise. ANOVA was used for comparisons among multiple groups. For survival analyses, statistical significance was assessed using the log-rank test. Correlations were evaluated using Spearman’s rank correlation, with the corresponding correlation coefficients and p-values reported.

---

**Figure S1.**
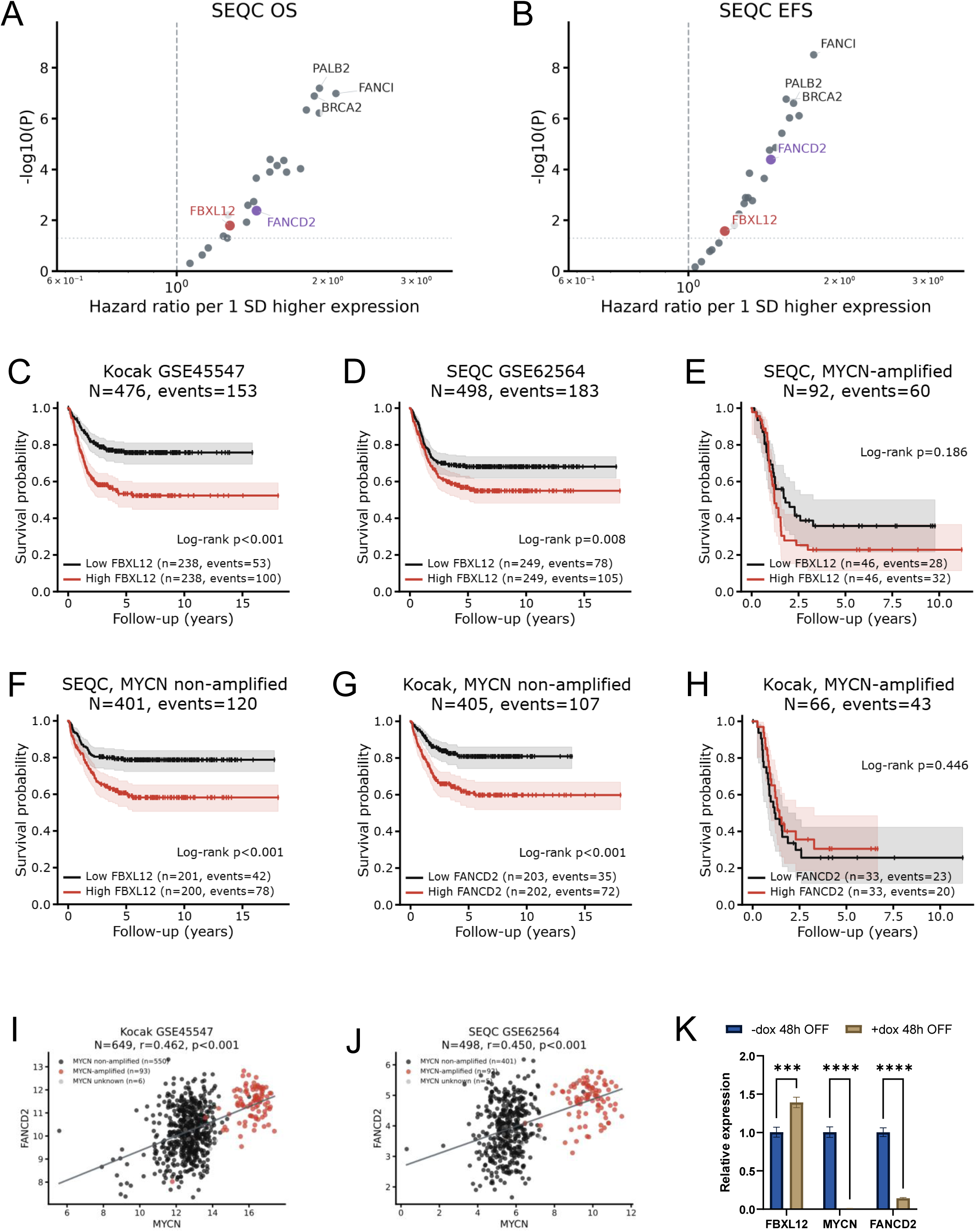
Extended clinical analyses of FBXL12, FANCD2 and MYCN-associated risk. (A, B) Univariate Cox proportional-hazards analysis of Fanconi anaemia pathway genes in the SEQC cohort for overall survival (A) and event-free survival (B). Hazard ratios are shown per 1 SD higher gene expression. FBXL12 and FANCD2 are highlighted. (C, D) Kaplan-Meier survival analysis stratified by median FBXL12 expression in the Kocak (C) and SEQC (D) cohorts. (E, F) Kaplan-Meier survival analysis stratified by median FBXL12 expression within MYCN-amplified (E) and MYCN non-amplified (F) SEQC cohort tumours. (G, H) Kaplan-Meier survival analysis stratified by median FANCD2 expression within MYCN non-amplified (G) and MYCN-amplified (H) Kocak cohort tumours. (I, J) Correlation between MYCN and FANCD2 mRNA expression in the Kocak (I) and SEQC (J) cohorts. Points are coloured by MYCN-amplification status. Pearson correlation was used for statistical analysis. (K) qPCR analysis of FBXL12, MYCN and FANCD2 mRNA expression in SHEP cells following doxycycline treatment for 48 h to suppress MYCN expression.

**Figure S2.**
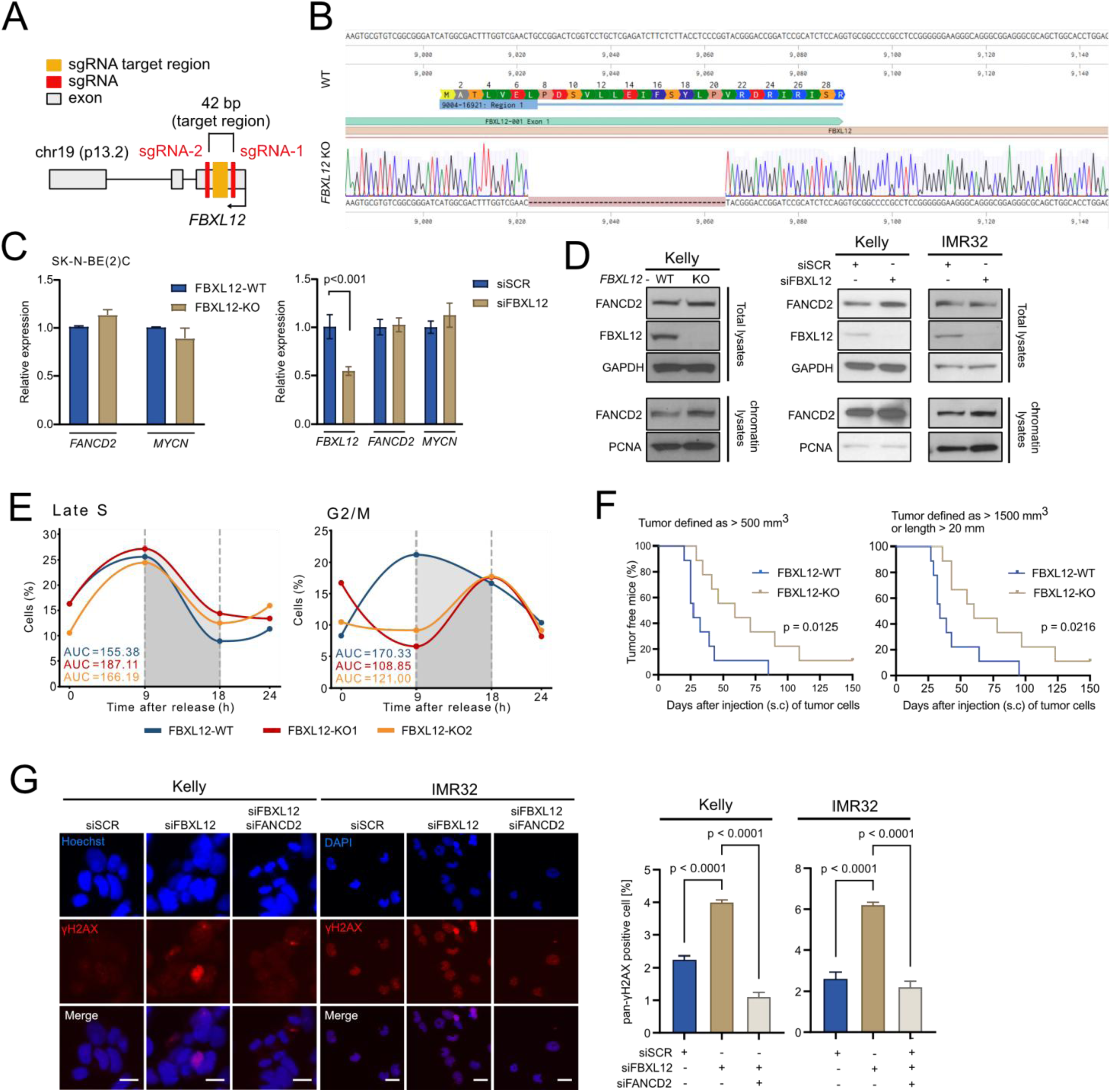
(A) Two sgRNAs designed to target the first exon of *FBXL12* and generate FBXL12-KO clones. (B) Validation of FBXL12-KO clones by Sanger-sequencing. (C) q-PCR analysis of FANCD2, MYCN and FBXL12 mRNA in SK-N-BE(2)C WT and FBXL12-KO clones or in SK-N-BE(2)C cells with FBXL12 depletion by siRNAs. (D) Left: Immunoblots of FANCD2 in total lysates or in chromatin lysates from Kelly WT and FBXL12-KO clones. Right: Immunoblots of FANCD2 in total lysates or in chromatin lysates from Kelly and IMR32 cells post 48-h transfection of FBXL12 siRNAs. (E) SK-N-BE(2)C WT and two FBXL12-KO clones were pulse-labelled with EdU for 30 min and released for the indicated times. EdU incorporation and DNA content (Hoechst, NN) were analysed by flow cytometry. The percentages of EdU-positive cells in late S phase and 4N cells in G2/M are shown over time. AUC was calculated over the 9-18 h interval (shown in grey). (F) Kaplan-Meier curves of the percentage of tumour free mice following subcutaneous (s.c) injection of SK-N-BE(2)C WT (blue, n = 9) and FBXL12-KO (light brown, n = 9) clones. Tumour formation was defined as a tumour volume greater than 500 mm^3^ (left panel) and 1500 mm^3^/length longer than 20 mm (right panel) with p-value of Logrank (Mantel-Cox) test. (G) γH2AX foci quantification upon FANCD2 rescue by FANCD2 siRNA co-depletion in siRNA-mediated FBXL12 knockdown IMR-32 and Kelly cell lines. (C and G) Data shown are means with error bars indicating SD.

**Figure S3.**
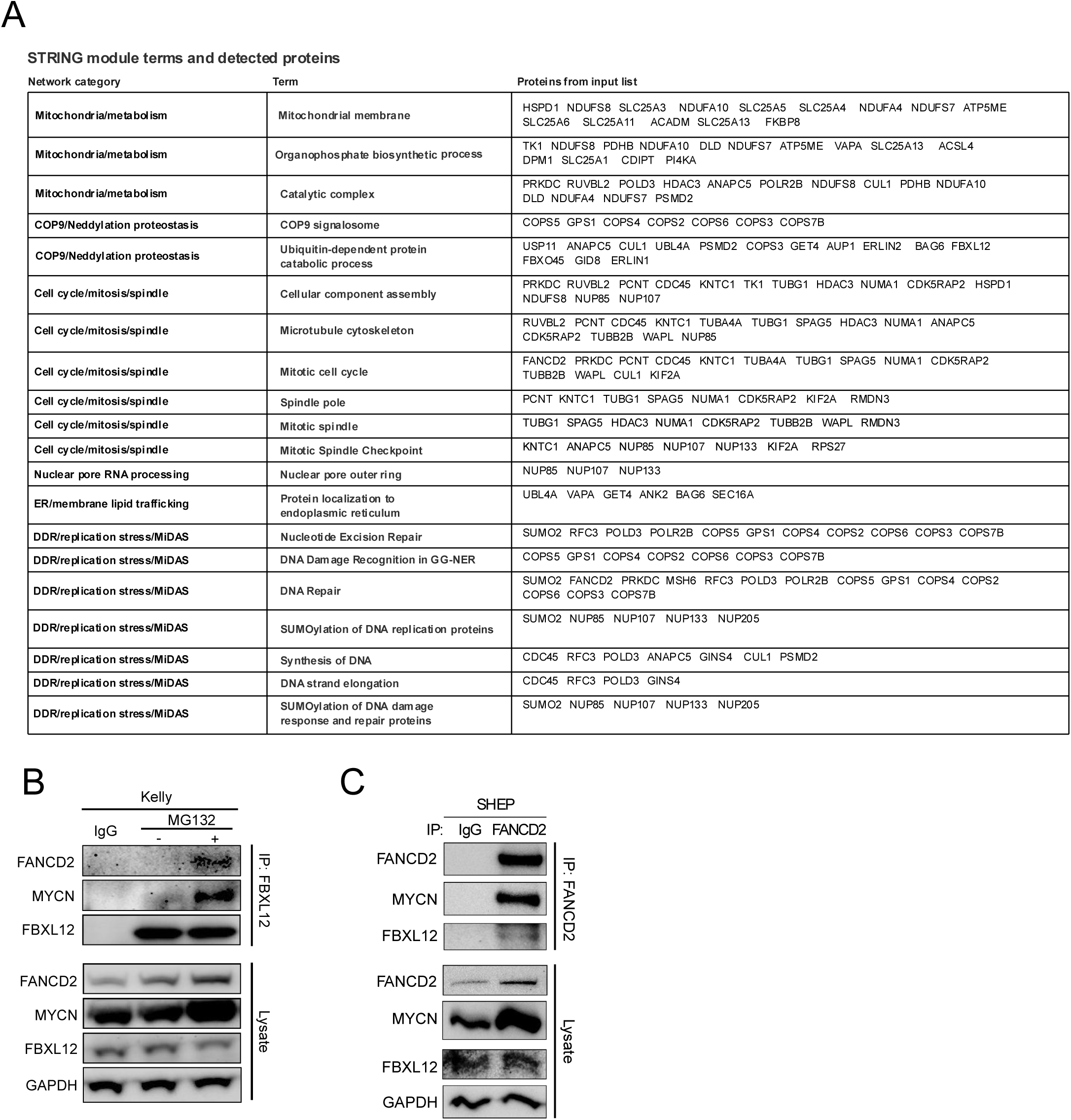
(A) Network categories correspond to the modules shown in the unbiased STRING protein interaction network in Figure 3B. Each row lists one enriched functional term and the input-list proteins contributing to that term. (B) Immunoprecipitation of endogenous FBXL12 from KELLY cell lysates following treatment with or without MG132 for 4h. (C) Immunoprecipitation of endogenous FBXL12 from MYCN-expressing SHEP cell lysates.

**Figure S4.**
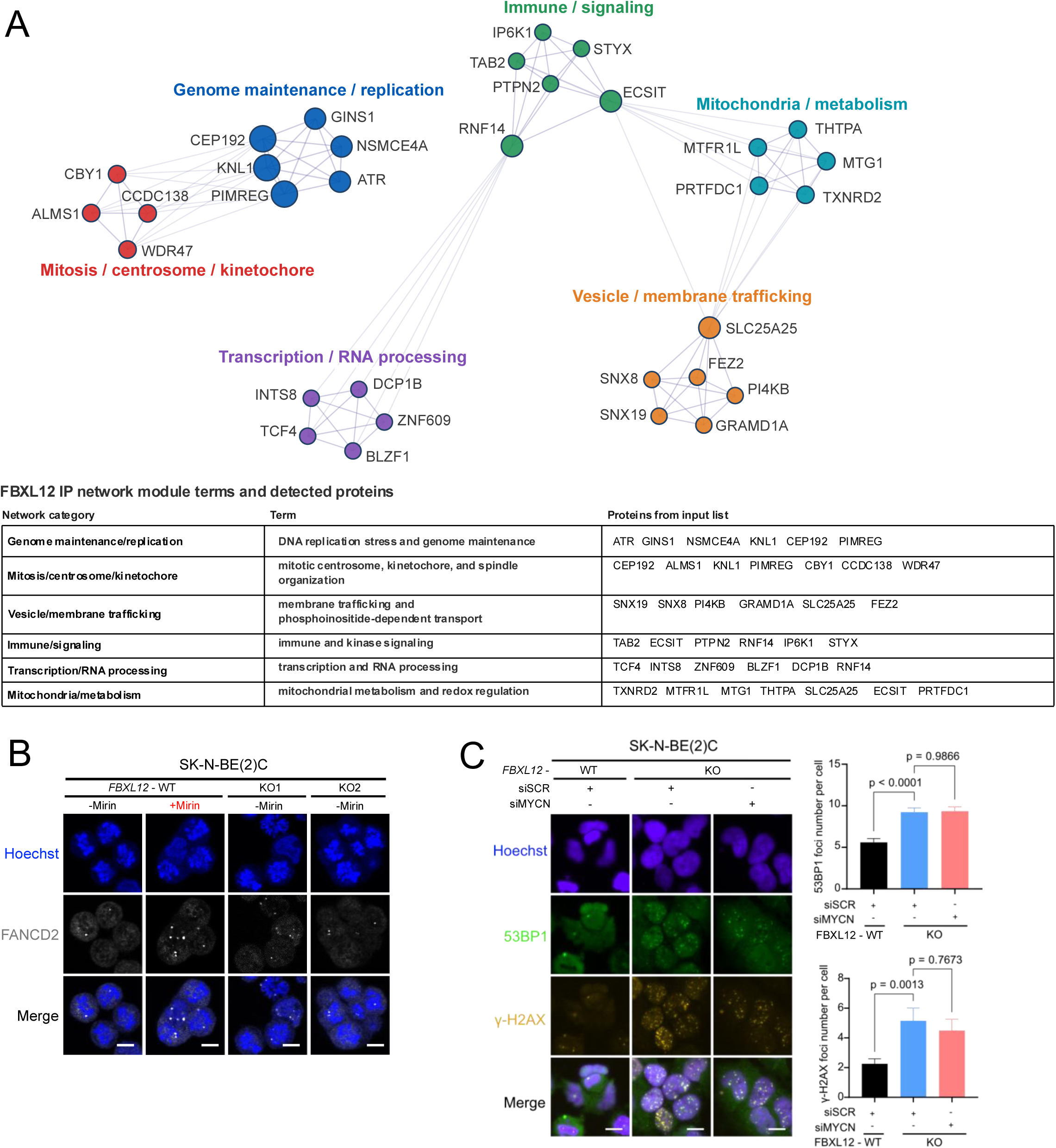
(A) Upper: Unbiased protein network generated from proteins identified in the FBXL12 immunoprecipitation mass spectrometry. Nodes represent FBXL12-associated proteins and edges indicate predicted functional associations. Node colours denote module assignment, and node size reflects interaction degree within the displayed network. Lower: Network categories correspond to the modules shown in the FBXL12 IP unbiased protein network. Each row lists the representative functional term assigned to each module. (B) Immunofluorescence of FANCD2 foci in SK-N-BE(2)C WT and FBXL12-KO cells at prometaphase, scale bar: 10 μm. (C) Quantification of 53BP1 and H2AX in SK-N-BE(2)C WT and FBXL12-KO G1 daughter cells with/without MYCN co-depletion, scale bar: 10 μm. Data shown in (C) are means with error bars indicating SD.

**Figure S5.**
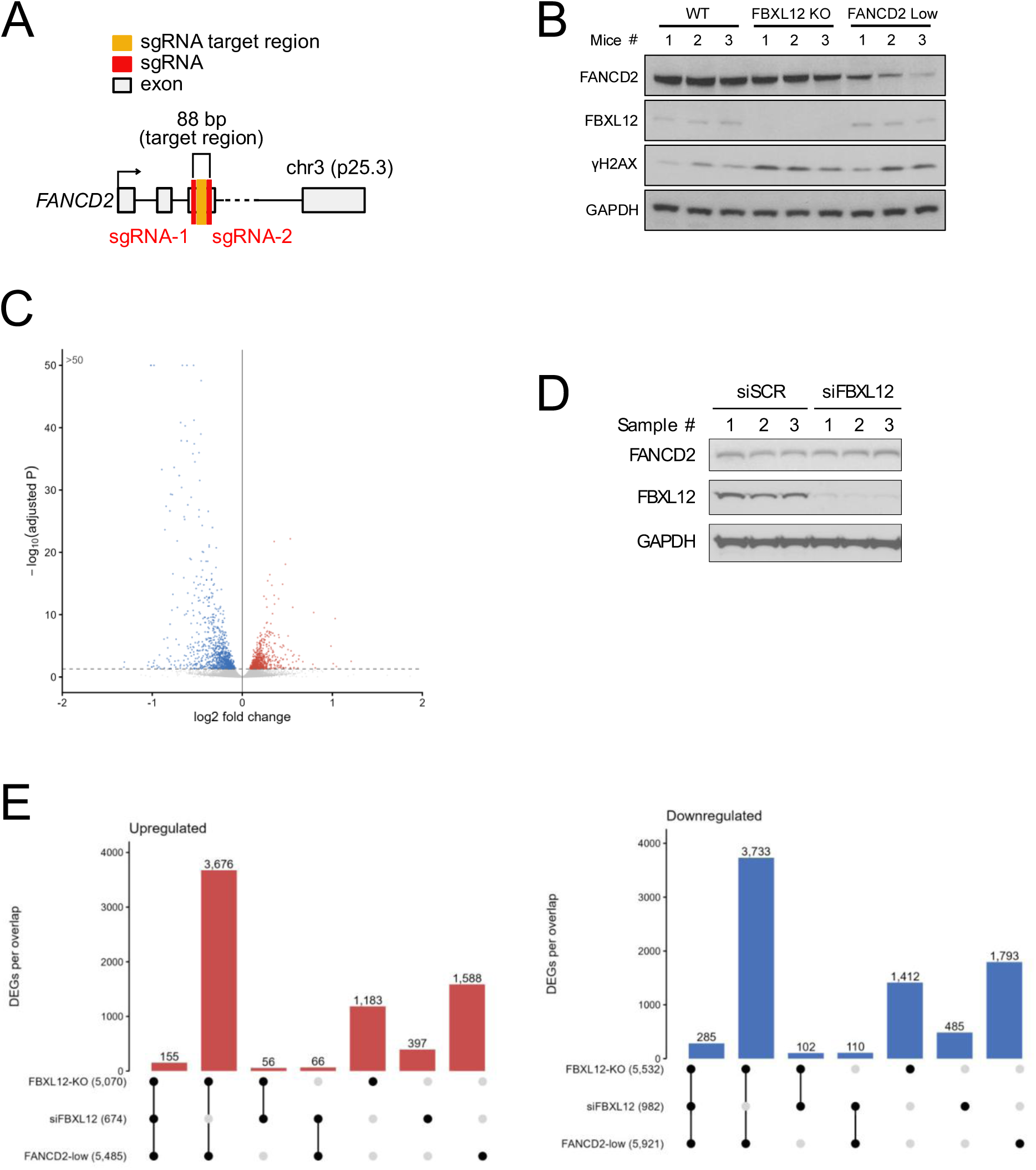
(A) Two sgRNAs designed to target the exon 3 of *FANCD2* and generate FANCD2-low clones. (B) Immunoblots of FBXL12, FANCD2 and γH2AX in total lysate of the resected tumours. (C) Volcano plot showing differential gene expression after acute FBXL12 depletion in SK-N-BE(2)C cells compared with siSCR control cells. (D) Immunoblot validation of siRNA-mediated target depletion in SK-N-BE(2)C cells used for RNA sequencing. Matched aliquots were processed for immunoblotting and RNA extraction 48 h after transfection. (E) Overlap of significantly upregulated and downregulated genes across FBXL12-knockout, siFBXL12-depleted and FANCD2-low cells. Bars indicate the number of genes in each intersection.

**Figure S6.**
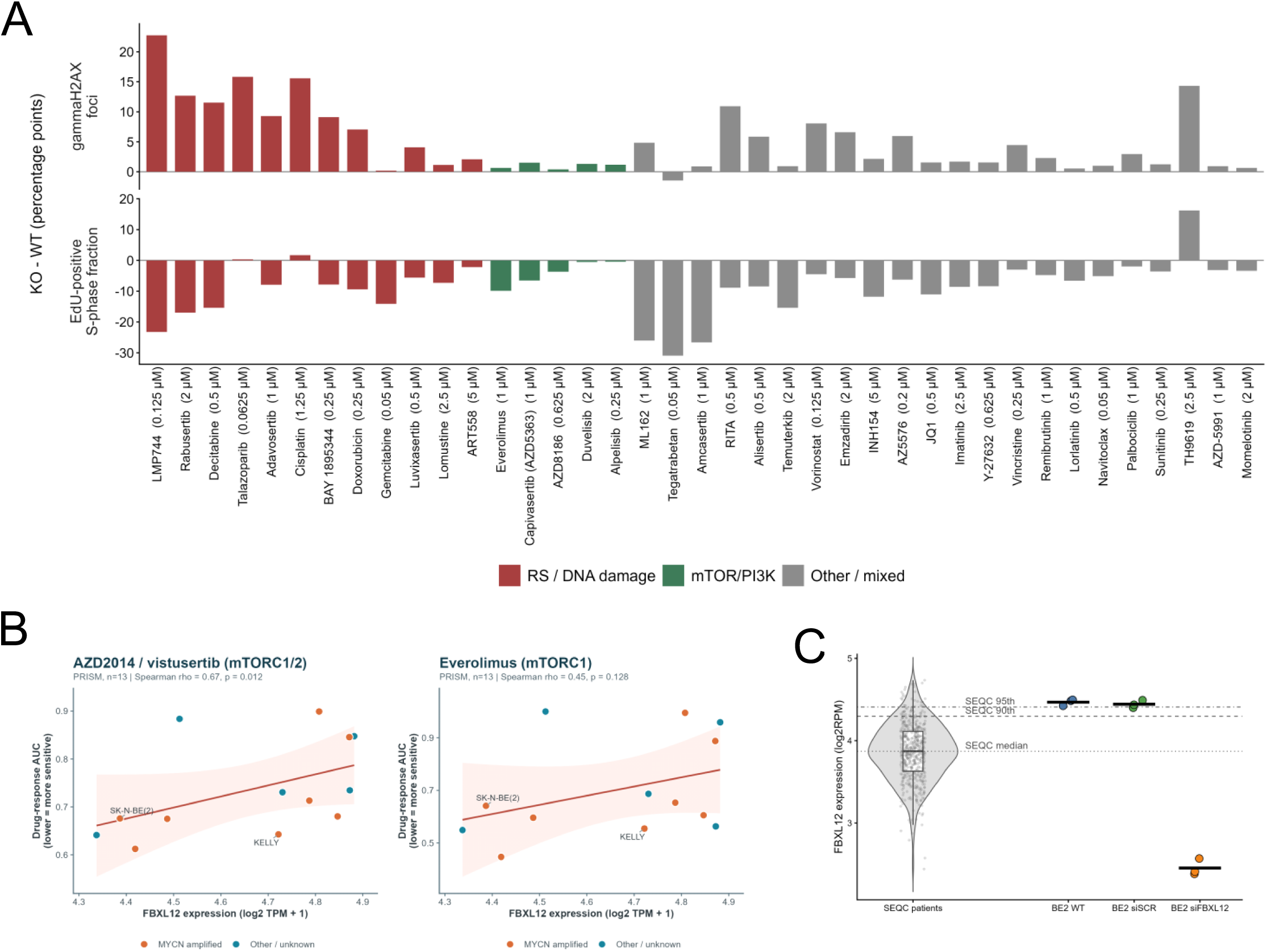
(A) Secondary focused high-content imaging drug screen in SK-N-BE(2)C wild-type and FBXL12-knockout cells. Bars show the mean knockout-minus-wild-type difference in the percentage of cells with >5 γH2AX foci (top) and the EdU-positive S-phase fraction (bottom). For each compound, the concentration shown was selected based on the strongest changes in γH2AX foci and S-phase occupancy. Positive values indicate higher values in knockout cells and negative values indicate lower values. Bars represent the mean of two replicate measurements; colours denote compound category. (B) Association between FBXL12 mRNA expression and PRISM drug-response AUC for AZD2014/vistusertib and everolimus in DepMap neuroblastoma cell lines. Each point represents one cell line; lower AUC indicates greater drug sensitivity. Points are coloured by MYCN-amplification status. The fitted trend line is shown, and Spearman correlation was used for statistical analysis. (C) Comparison of FBXL12 expression in SEQC neuroblastoma patient tumours and SK-N-BE(2)C RNA-seq samples. Patient tumours are shown as a distribution, with BE(2)C wild-type, siSCR and siFBXL12 samples overlaid separately.

